# Long-distance gene flow and recombination shape the evolutionary history of a maize pathogen

**DOI:** 10.1101/2024.07.22.604531

**Authors:** Flávia Rogério, Cock Van Oosterhout, Stéphane De Mita, Francisco Borja Cuevas-Fernández, Pablo García-Rodríguez, Sioly Becerra, Silvia Gutiérrez-Sánchez, Andrés G. Jacquat, Wagner Bettiol, Guilherme Kenichi Hosaka, Sofia B. Ulla, Jürg Hiltbrunner, Rogelio Santiago, Pedro Revilla, José S. Dambolena, José L. Vicente-Villardón, Ivica Buhiniček, Serenella A. Sukno, Michael R. Thon

**Affiliations:** University of Salamanca, Department of Microbiology and Genetics, Institute for Agribiotechnology Research (CIALE), Villamayor, Salamanca, Spain; School of Environmental Sciences, University of East Anglia, Norwich Research Park, Norwich, UK; PHIM Plant Health Institute, Univ Montpellier, INRAE, CIRAD, Institut Agro, IRD, Montpellier, France; Faculty of Exact, Physical and Natural Science, National University of Córdoba, IMBIV-CONICET-ICTA, Córdoba, Argentina; Embrapa Environment, Jaguariúna, São Paulo, Brazil; Laboratory of Genetics of Microorganisms “Prof. Joao Lucio de Azevedo”, Department of Genetics, “Luiz de Queiroz” College of Agriculture, University of São Paulo, Piracicaba, Brazil; Federal Department of Economic Affairs, Agroscope, Centre of competences Plants and Plant Products, Zurich, Switzerland; Misión Biológica de Galicia, Spanish National Research Council (CSIC), Pontevedra, Spain; Statistics Department, University of Salamanca, Salamanca, Spain; Bc Institute for Breeding and Production of Field Crops, Dugo Selo, Croatia

**Keywords:** *Colletotrichum graminicola*, population genomics, restriction site-associated DNA sequencing (RAD-seq), whole-genome sequencing (WGS), isolation by distance (IBD), genetic introgression

## Abstract

The evolutionary history of crop pathogens is shaped by a complex interaction of natural and anthropogenic factors. The ascomycete fungus *Colletotrichum graminicola* causes maize anthracnose. The disease can result in significant yield losses and is also an important model for genetic studies. We conducted a comprehensive investigation into the evolutionary genomics of *C. graminicola* using a collection of 212 isolates from 17 countries. Genomic analyses supported the existence of three geographically isolated genetic lineages, with a significant pattern of isolation by distance. We identified two distinct gene flow patterns, driven by short and long-distance dispersion, likely resulting from the natural spread of the pathogen and the exchange of contaminated seeds. We present evidence of genetic introgression between lineages, suggesting a long history of recombination. We identified significant recombination events coalescing at distinct points in time, with the North American lineage displaying evidence of most ancient recombination. Demographic modeling indicated that North America is an intermediate between Brazil, Europe and an ancestral, unsampled source population, hypothesized to be Mesoamerican. Our analyses revealed that the global genomic structure of *C. graminicola* is shaped by geographic differentiation driven by long-distance migration and a long history of recombination and introgression. We show historical relationships among these lineages, identifying a potential route for fungal spread, with the North American population emerging ancestrally, followed sequentially by the Brazilian and European populations. Our research indicates that the European lineage is more virulent, which has implications for the potential emergence of new outbreaks of maize anthracnose in Europe.

## Introduction

Examining the processes that drive species evolution over time is a central focus of evolutionary genomics. Molecular population genetics provides valuable insights into patterns of genetic variation within populations, enhancing our comprehension of their evolutionary history and population dynamics (Dobzhansky 1973; Casillas and Barbadilla 2017; Gladieux et al. 2011). This knowledge has practical applications, for instance for studying how populations of plant pathogens will respond to future environmental changes and selective pressure, or for determining the role human activities play in the dispersion of pathogens, which is crucial for implementing effective disease control measures (McDonald and Linde 2002; Giraud et al. 2008; Everhart et al. 2021).

The genetic composition of pathogen populations is shaped by evolutionary forces and interactions with hosts and local environments (Charlesworth 2010). Anthropogenic changes to the environment have a large impact on the interactions between the evolutionary forces, which in turn have altered the Red Queen Dynamics between parasites and their hosts (Van Oosterhout 2021). Migration encompasses individual or propagule movement, leading to gene flow between isolated gene pools (McDermott and McDonald 1993). Understanding migration patterns is vital for informed disease prevention decisions, as it may introduce new genetic variations, including virulence factors, and connect adapted populations (McDonald and Linde 2002). Together with gene flow, genetic recombination constitutes vital evolutionary forces driving variation and pathogen adaptation (Milgroom 2015; de Vries et al. 2020; Stukenbrock and Bataillon 2012). Genetic recombination has the potential to accelerate adaptation by the generation of new, potentially adaptive genotypes that were favored by natural selection, or combining adaptive alleles at distinct loci that appeared in different genetic backgrounds (Barton 2010). This process offers new substrates for adaptive evolution while simultaneously purging the genome of deleterious mutations accumulated over time (Van Oosterhout 2021; Taylor et al. 1999; Felsenstein 1974).

The fungus *Colletotrichum graminicola* is an important maize pathogen, associated with anthracnose disease and responsible for significant yield losses (Bergstrom and Nicholson 1999; Frey et al. 2011; Sukno et al. 2008). This haploid ascomycete belongs to the Graminicola species complex, a well-defined monophyletic clade comprising species mainly associated with grass hosts (Talhinhas and Baroncelli 2021; Bhunjun et al. 2021; Talhinhas and Baroncelli 2023). It caused devastating outbreaks in the USA during the 1970s and persists as an established endemic disease, largely managed through resistant cultivars (Byrnes and Carroll 1986; Lipps 1983; Rogério et al., 2023). Recent reports of maize anthracnose occurrences in multiple countries imply that the geographic range of the pathogen is expanding (Cuevas-Fernández et al. 2019; Rogério et al. 2023b). Considering its historical impact and the ongoing influence of climate changes on plant disease dynamics, anthracnose could pose an increasingly severe threat to maize cultivation worldwide.

In addition to its economic importance, *C. graminicola* is also an important model for studying plant-pathogen interactions, being the pioneer among *Colletotrichum* species to undergo genome sequencing (O’Connell et al. 2012; Perfect et al. 1999; Becerra et al. 2023). The genetic structure of *C. graminicola* has been explored globally (Rogério et al. 2023). Population genomics analyses have revealed that the pathogen is structured into three largely isolated populations associated with North America, Europe, and Brazil. Genetic exchanges between these populations are facilitated by intra- and intercontinental migration, with evidence of genetic recombination. Nevertheless, the current impact of gene flow and recombination at the genome level, as well as the evolutionary origin of these populations, remains unknown.

Population genomics studies of many crop pathogens demonstrated that human activities (domestication, trade, and migration) have played a pivotal role in the emergence and spread of plant diseases (Goss et al. 2014; Chakraborty 2013; Tabima et al. 2020; Kamvar et al. 2015b; McMullan et al. 2018; Sotiropoulos et al. 2022; Barrès et al. 2008). The dissemination of new virulence genes and agrochemical resistance among populations across large geographical areas is significantly influenced by gene flow, which plays a central role in driving pathogen evolution (Zhan et al. 2015). Like mutation and recombination, gene flow is a source of new genetic variation, which forms the substrate for selection in host-parasite interaction. By introducing novel alleles from other populations, gene flow indirectly increases the effective population sizes of the pathogen. The increased rates of gene flow in our human-modified environment give present-day pathogens an important coevolutionary advantage (Van Oosterhout 2021). Understanding and monitoring gene flow is crucial for developing effective strategies to manage and mitigate the potential risks associated with that. Competition theory predicts that when less related genotypes compete, it may favor the selection of more aggressive strains (Koskella et al. 2006; Zhan and McDonald 2013; López-Villavicencio et al. 2007). In this context, gene flow between genetically divergent lineages can decrease genetic relatedness and potentially promote the emergence of these more aggressive strains. This can pose a risk factor for host resistance breakdown.

Fungi exhibit a range of reproductive strategies, involving genetic recombination outside conventional sexual reproduction, such as hyphal fusion (anastomosis) and parasexual cycles (Giraud et al. 2008). While clonality is the predominant mode of reproduction, population genetic studies highlight that recombination plays a key role in the genetic structure of phytopathogenic fungi (Taylor et al. 2015; Stukenbrock and Dutheil 2018; Meng et al. 2015; Drenth et al. 2019; Ashu et al. 2017; Roca et al. 2005). Notably, recombination is reported to be an important factor in the ecology and evolutionary dynamics of numerous *Colletotrichum* species (Rogério et al. 2022; Souza-Paccola et al. 2003; Souza et al. 2010). However, certain studies have elucidated alternative mechanisms, such as the parasexual cycle, contributing to genetic recombination in *Colletotrichum* genus (Souza-Paccola et al. 2003; Rosada et al. 2010; Barcelos et al. 2014; Nordzieke et al. 2019).

Gene flow and recombination lead to genomes with admixed ancestry, a process that is commonly referred to as genetic admixture (Beichman et al. 2018). The genomic signature of admixture is characterized by chromosomal segments that originate from distinct genetic backgrounds or populations (Carroll 2013). Closely affiliated to admixture is the term “introgression”. Introgression is a process resulting from recombination and gene flow between significantly diverged gene pools, leading to the gradual transfer of genetic material from one pool to the other through admixed individuals (Anderson 1953; Stukenbrock 2016). At the far end of this continuum is horizontal gene transfer, which is the genetic exchange between otherwise reproductively isolated species. Here in this study, we have adopted the term introgression to refer to genetic exchanges between divergent lineages. These genetic exchanges leave distinct signatures in the genome, identified through DNA sequence comparisons, with introgressed regions typically exhibiting high nucleotide similarity, followed by a sudden change in divergence (Gompert and Buerkle 2013; Ward and van Oosterhout 2016; Rosenzweig et al. 2016).

The availability of genomic data enables the determination of the genomic ancestry of fungal pathogens, shedding light on their evolutionary biology. For instance, the generalist parasite *Albugo candida* has 25% of its genome introgressed among physiological races, thereby facilitating the exchange of effector repertoires (McMullan et al. 2015). In *C. truncatum*, recent introgression between coexisting lineages was detected, involving protein-encoding genes, suggesting potential co-evolutionary targets for selection (Rogério et al. 2022). Likewise, *Cryptosporidium parvum*, a zoonotic fungus causing disease in both humans and ruminants, multiple recent genetic exchanges were identified between lineages, affecting about 22% of the genome, with introgressed regions significantly enriched with putative virulence genes (Corsi et al. 2022).

In this study, we focused on unraveling the evolutionary dynamics shaping the genetic structure of *C. graminicola* by using genomic data. Our investigation employed a global collection of 212 isolates from 17 countries, allowing for a comprehensive analysis of genetic exchanges on a genome scale and determination of the evolutionary origin of the *C. graminicola* lineages. We also explored the impact of isolation by distance and identified gene flow patterns responsible for the population structuring, and we investigated the role of genetic introgression between genetically divergent lineages. Finally, through demographic modeling, we unveiled the historical relationship among lineages and identified a potential route for the expansion of the fungus from its initial source population. By providing a comprehensive analysis of these genetic mechanisms, our study contributes to a deeper understanding of the evolutionary genomics of *C. graminicola*, with implications for disease management.

## Material and Methods

### Data processing

In total 212 isolates of *Colletotrichum graminicola* were obtained from field samples and public culture collections from seventeen countries (Table S1). Mycelia from monosporic cultures were incubated on an orbital shaker in potato dextrose broth medium (PDB) and genomic DNA was extracted using DNeasy Plant Mini Kits (Qiagen Inc., Valencia, California, USA). Ninety-four isolates were obtained from Rogério et al. (2023), and 118 isolates were newly isolated and sequenced, using restriction site-associated DNA sequencing (RAD-seq) or whole-genome sequencing (WGS). New 82 isolates were genotyped with RAD-seq by Floragenex Inc. (Beaverton, OR, USA). RAD-seq libraries with sample-specific barcode sequences were constructed from DNA digested with restriction enzyme PstI, 202 then single- end sequenced (1 x 100 bp) on one lane of an Illumina HiSeq 2000 instrument (San Diego, CA), by (Beaverton, OR, USA). Whole genome resequencing of 36 isolates was performed by BGI company (Hong Kong, China) using the DNBSEQ Libraries sequencing platform (2 x 150 bp).

The RAD-seq reads were demultiplexed and quality filtered using the PROCESS_RADTAGS module of the software STACKS v.2.10 (Rochette et al. 2019). The demultiplexed reads were aligned to the *C. graminicola* reference genome M1.001 (NCBI accession number PRJNA900520) using BWA v.0.7.8 (Li and Durbin 2009). For the WGS reads, the sequence quality was checked with FASTQC v0.11.7 (Babraham Bioinformatics), and low-quality reads were trimmed with TRIMMOMATIC v0.3.8 (Bolger et al. 2014). Alignments were sorted and each bam file was assigned to a read group with SAMTOOLS v. 1.3 (Li et al. 2009). Variant calling was performed with the HaplotypeCaller module of GATK v.3.7, applying hard filters as suggested in GATK’s Best Practices (Van der Auwera et al. 2013). Additional filters were applied using VCFTOOLS (Danecek et al. 2011). Only biallelic SNPs with a genotype rate of 90% and sites with a minor allele frequency of less than 5% were retained.

### Global population structure

The presence of repeated multilocus genotypes (here referred to as clones) was identified using the function MLG.FILTER with a threshold determined by the CUTOFF_PREDICTOR tool based on Euclidean distance using the package POPPR v2.8.6 (Kamvar et al. 2015). Population genetic analyses were conducted on a clone-corrected dataset (i.e., only one representative of each repeated multilocus genotype was retained). To investigate the populational subdivision, first, we used the clustering method implemented in the SNMF software to infer individual ancestry coefficients, which is more appropriate to deal with inbred or clonal lineages because it does not assume Hardy-Weinberg equilibrium (Frichot et al. 2014). The best number of clusters (*K*) was chosen using the cross-entropy criterion based on the prediction of masked haplotypes to evaluate the error of ancestry estimation and the quality of fit of the model to the data. We ran 10 repetitions for each value of *K* ranging from 1 to 10. To confirm the populational differentiation we employed a principal component analysis (PCA), with *a priori* geographical knowledge from sampling (by continent) and a discriminate analysis of principal components (DAPC), both analyses conducted in the R environment with the package ADEGENET v2.0.1 (Jombart and Ahmed 2011). The FIND.CLUSTER function was used to identify the most likely grouping in the data based on the Bayesian information criterion (BIC) calculated for 1 to 10 clusters. Additionally, we reconstructed a neighbor network using the software SPLITSTREE v4 (Steinrücken et al. 2019) to visualize phylogenetic signals based on a distance matrix with default parameters.

To further understand the impact the geographic subdivision on the population structure we conducted isolation-by-distance analyses. We computed pairwise genetic dissimilarity and geographical distance between samples and examined their relationship using regression analysis. The correlation between the matrices of genetic and geographic distance was also observed using a simple Mantel test with 1000 permutations as implemented in the VEGAN package in R (v.2.6.4). Genome variability was assessed through nucleotide diversity statistics computed using EGGLIB v.3.3.2 (Siol et al. 2022).

### Footprints of recombination

To examine the impact of genetic exchanges on tree topologies we constructed a consensus tree using DENSITREE 2 software (Bouckaert 2010). We used BEAUTI software v1.5.3 to generate XML files for each contig. Substitution models were selected from BMODELTEST (Bouckaert and Drummond 2017), which detects the best model and automatically adds it to the XML file. BEAST 2 (v2.5.1) (Bouckaert et al. 2019) was used to generate phylogenetic trees using the Markov Chain Monte Carlo (MCMC) for 1×10^6^ generations with sample frequency of 2,000 and burn-in of 10%.

Recombination was analyzed in multiple genome alignments by chromosome including non-variant sites. We used the module GENOTYPEGVCS in GATK to include genotypes from all positions present in the reference genome and BCFTOOLS v.10.2 (Li et al. 2009) to obtain consensus sequences for each chromosome. Recombination events were analyzed by the RDP4 software (Martin et al. 2021) using seven independent algorithms (RDP, GENECONV, BOOTSCAN, MAXCHI, CHIMAERA, SISCAN, AND 3SEQ). Significant events (*P*<0.05) were considered if they were significant with three or more detection methods (Jouet et al. 2015). Only events for which the software identified the parental sequences (i.e., no “unknowns”) were considered. To visualize the significant recombinant blocks identified by RDP4, we used the HYBRIDCHECK software (Ward and van Oosterhout 2016) with a step size of 1 and a window size of 500 SNPs. The HYBRIDCHECK software was also used to date the recombination events. The divergence times between the donor and recombinant sequence were calculated using a JC69 mutation correction, assuming a mutation rate of 10^−8^ per generation and a generation time of one year (Kasuga et al. 2002).

Recombination events were also dated through Bayesian inference in BEAST 2. We selected some recombinant blocks detected by HYBRIDCHECK and applied a Bayesian coalescent approach for dating. XML files were created using BEAUTI, where the module BMODELTEST was used to determine the optimal substitution model under a mutation rate of 10^−8^ and strict clock rate. We assumed the model of Coalescent Constant Population with default priors and chain length of 1×10^7^ generations. Each dating analysis was replicated five times. TRACE was used for visualizing posterior parameters. The Treeheight statistic, representing the marginal posterior distribution of the age of the root of the entire tree, was considered as the divergence time from the common ancestor (tMRCA) for the set of sequences evaluated.

For dating, we assumed a generation time of 1 year. Note that the reported dates are likely an overestimate of the true age of recombination. In other words, the “true” recombination event may be more recent than our inferred estimate. There are two ways in which the dating estimates can be biased, and both lead to an overestimation of the true date of introgression. First, there is an overestimation because the descendants of the actual donor and recipient sequences are unlikely to have been sampled. Consequently, the introgressed regions look more diverged (and thus “older”). Second, even if the direct descendants of the donor sequence were included in our analysis, they may have diverged from the block in the recipient block due to subsequent recombination. This may have introduced multiple nucleotide polymorphisms wholesale into the introgressed region. Given that we assume that divergence is driven by the mutation rate (i.e., a molecular clock), polymorphisms introduced by recombination would make the divergence appear older (see Jouet et al. 2015). Both processes lead to an overestimation of the true date of introgression.

### Demographic inference

We inferred the evolutionary history of *Colletotrichum graminicola* genetic lineages using Approximate Bayesian Computation (ABC) based on random forest (RF), conducted in the Python package PYABCRANGER (Collin, et al. 2020). We based our analysis on three genetic lineages (North American, Brazilian, and European) treated as independent populations. We used the Python package EGGLIB to perform coalescent simulations and to compute the summary statistics, and PYABCRANGER to perform model choice and parameter estimation. ABC-RF uses a classification vote system after bagging (i.e., aggregating bootstrap results) the simulated outputs to identify the most probable model. The model with the highest number of votes is deemed to be the most probable model. To shed light on divergence events and colonization routes, we modeled 22 scenarios of divergence including asymmetrical migration and bottleneck events, assuming either stepping-stone and single-source models. Models with and without an unknown population (i.e., ‘ghost’ population) as an ancestral population were tested. In our models, every divergence event was immediately followed by a bottleneck, otherwise keeping constant population size for all populations. For detailed methodology, see Supplementary text S1.

### Pathogenicity assays

Pathogenicity assays were performed by *in vivo* leaf blight assays, following Vargas et al. 2012), as described by Rogério et al. (2023). A total of 56 isolates were evaluated, and distributed across batches due to the limitations of accommodating all isolates simultaneously during the pathogenicity assays. Recognizing the potential for a batch effect, we selected the reference strain M1.001 to be included in all batches, enabling the estimation of systematic differences among them. This strain was originally collected from North America in 1972 during the US outbreak and is highly virulent (Rogério et al. 2023; O’Connell et al. 2012). To assess batch effects, we compared the repeated analysis of M1.001 strain across different batches using an analysis of variance (ANOVA), with lesion size as the response variable, batch as the predictor variable, and a null hypothesis of no difference between batches. The obtained highly significant differences (*P*<0.0001), indicated a clear batch effect. Consequently, to mitigate these systematic differences among batches, we standardized each strain using the mean and standard deviation of the control strain specific to its batch. An ANOVA of the standardized data was then employed to identify significant differences among isolates. Subsequently, a second test aimed at determining differences among isolates while accounting for batch variation in lesion sizes. The predictor variable was isolate, the response variable was the corrected lesion size, and the null hypothesis assumed no difference in lesion size among isolates. The ANOVA test was significant (*P*<0.0001), so we performed a Sidäk’s comparison-of-means test. This choice was made because it is exact for independent comparisons and easily translated into a graphical representation, facilitating the identification of significant differences among strains. Importantly, Sidäk’s test is less restrictive than the commonly used Bonferroni test for these types of problems.

The isolates that had been tested in the virulence assay were divided into groups based on sampling year and genetic structure. The isolates were classified into three categories based on genetic group and virulence (less, equal, or more virulent than isolate M1.001) and subjected to a Fisher’s exact test. The isolates were also grouped by year into three categories (before 2012, between 2012 and 2016, and after 2016), and subjected to a Kruskal–Wallis test. The statistical tests and figures were implemented in the statistical programming language R.

Finally, we examined whether there was a positive correlation between pairwise genetic distance between isolates and similarity in their level of virulence. Genetic distance was calculated as the difference in the number of bases in pairwise comparisons. The similarity in virulence level was assessed by comparing the difference between lesion areas (calculated as the lesion area of isolate A minus the lesion area of isolate B). The correlation between pairwise genetic distance and similarity in virulence was analyzed using a Mantel test with 1000 permutations as implemented in the VEGAN package in R (v.2.6.4).

## Results

### Population structuring

We characterized the global genetic structure of *C. graminicola* using genomic data from 212 isolates based on 7,207 biallelic single nucleotide polymorphisms (SNPs) distributed across 13 chromosomes, with an average of 1.3 SNPs per 10kb (Fig. S1; Table S1). After clone correction, 166 unique multilocus genotypes were kept in the dataset (Table S2). A neighbor-net network revealed three distinct genetic groups, with isolates largely grouped by their geographic origin: South America, Europe (including Argentina), and North America (Fig. 1; Fig. S2). The observed population subdivision corroborates the conclusion drawn in our previous study (Rogério et al. 2023), which was based on a smaller number of isolates (Fig 1. 108 versus 212 presently). By including more genetic variants we obtained a better resolution of the genetic diversity distribution, but clustering methods still supported the existence of these groups. Henceforth, the three groups will be referred to as Brazilian (BR), European (EU), and North American (NA) lineages, respectively.

**Figure 1.**
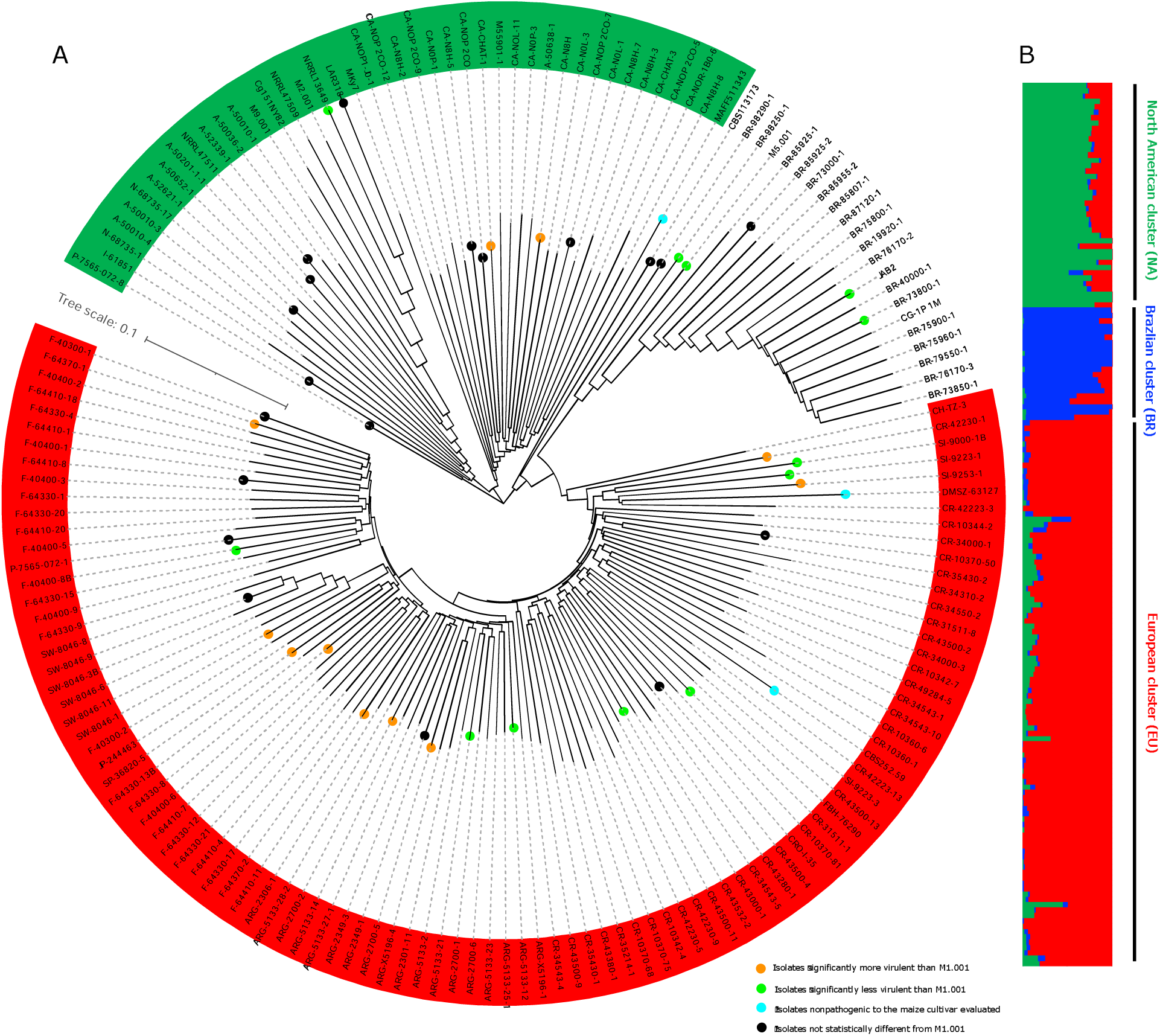
Population subdivision of *Colletotrichum graminicola* based on 7,207 single-nucleotide polymorphisms on the clone-corrected dataset. (A) Neighbor-net network showing relationships between isolates, visualized in a circular tree using iTOL v. 6 (Letunic and Bork 2007). Dots represent isolates with pathogenicity essayed. Samples SW-8046-2, P-7565-072-6, SW-8046-13, CR-49298-1, SL-9253-4, CRO-I-41, P-7565-072-3, CR-10342-5, F-64330-7, and F-64330-2 were removed from the network because they are clones. (B) Individual ancestry proportions in *K* = 3 clusters estimated with SNMF; each genotype is represented by a vertical bar.

These lineages exhibit different levels of overall genetic diversity, as indicated by estimated genomic indices, suggesting independent evolutionary trajectories (Table S3). The North American lineage possesses the highest estimates of genetic variation. Genetic diversity can be increased by a multitude of factors, including demographic effects. We formulated four (non-exclusive) hypotheses explaining the larger level of diversity in the NA lineage: 1) NA has a larger contemporary effective population size, or (2) NA is the ancestral (source) population, and other populations experienced a bottleneck when they emerged. (3) NA is experiencing recurrent incoming gene flow, and/or (4) NA has higher levels of recombination (4) NA has higher levels of (sexual or parasexual) recombination, limiting the consequences of background selection. Below we examine these hypotheses in more detail.

Population substructuring and admixture were analyzed in further detail using the non-parametric method implemented in SNMF. Based on the cross-entropy criterion, which was confirmed by the Bayesian information criterion, the model with *K* = 3 clusters best captures the population substructing in the whole dataset (Fig. S2). In addition, many genomes show evidence of admixture, which suggests genetic exchanges between *C. graminicola* genetic lineages (Fig. 1b; Table S4). Some isolates are highly admixed, for instance, P-7565-072-8, MAFF511343, CBS113173, M5.001, BR-98290-1, and BR-98250-1. These isolates are responsible for the “loops” or reticulate nodes in the neighbor network (Fig. S3). In contrast, a small proportion of isolates are completely pure i.e., 20% of isolates have individual ancestry coefficients that assign them to a single cluster (*q*>0.99, where *q* represents the proportion of an individual genome originating from a given ancestral gene pool) and are considered pure (Fig. S3, Table S4). The Brazilian cluster showed the largest proportion of pure isolates. North American and Brazilian lineages showed the most ancestry shared with Europe, with isolates showing membership lower than 1% with other clusters (Table S4). There was one exception, i.e., the isolate MAFF511343, which originated from a public culture collection. Interestingly, older North American isolates are genetically pure: LAR318 (1990), M2.001 (1975), Cg151NY82 (1982), M9.001 (1990), NRLL13649 (1988), NRLL47509 (2005). In addition, the NA lineage has the deepest rooting branch, and these isolates have the longest terminal branches, indicating a more ancient origin of this group.

The European clade encompasses all samples from European countries, one isolate from China and Japan, and all isolates from Argentina (see highlighted samples in Fig. S3). Except for some Portuguese samples (P-7565-072-4, P-7565-072-6, P-7565-072-7, P-7565-072-8), all isolates were grouped with samples from the same country of origin. To explore the genetic diversity in the European cluster we performed an analysis of molecular variance (AMOVA), with the function AMOVA from the POPPR package, using isolates from each country as independent populations. We excluded samples from countries with few representative isolates (i.e., samples CBS252.59, DMSZ-63127, FBH-76290, P-7565-072-1, P-7565-072-8, SP-36820-5, and CH-TZ-3). The AMOVA test revealed that 86% of genetic variation was within countries and 14% was among them (*P*<0.01). Clustering analyses showed a weak genetic differentiation within this lineage with isolates grouped mostly according to their countries of origin (Fig. S4). Although the methods employed for population subdivision did not show a clear number of subgroups, a neighbor-net network inferred with SPLITSTREE and PCA showed distinct subgroups grouped according to their country of origin, except for Croatia and Slovenia (Fig. 2). Isolates from Slovenia were localized close to Croatia, consistent with the close geographical proximity of both countries. However, the isolates SI-9000-1B and SI-9223-1 are within the Croatia subgroup, but apart from another sample SI-9223-3, suggesting independent introductions or gene flow between these countries. Interestingly, the samples from Argentina form a distinct group within European countries, suggesting an independent introduction event.

**Figure 2.**
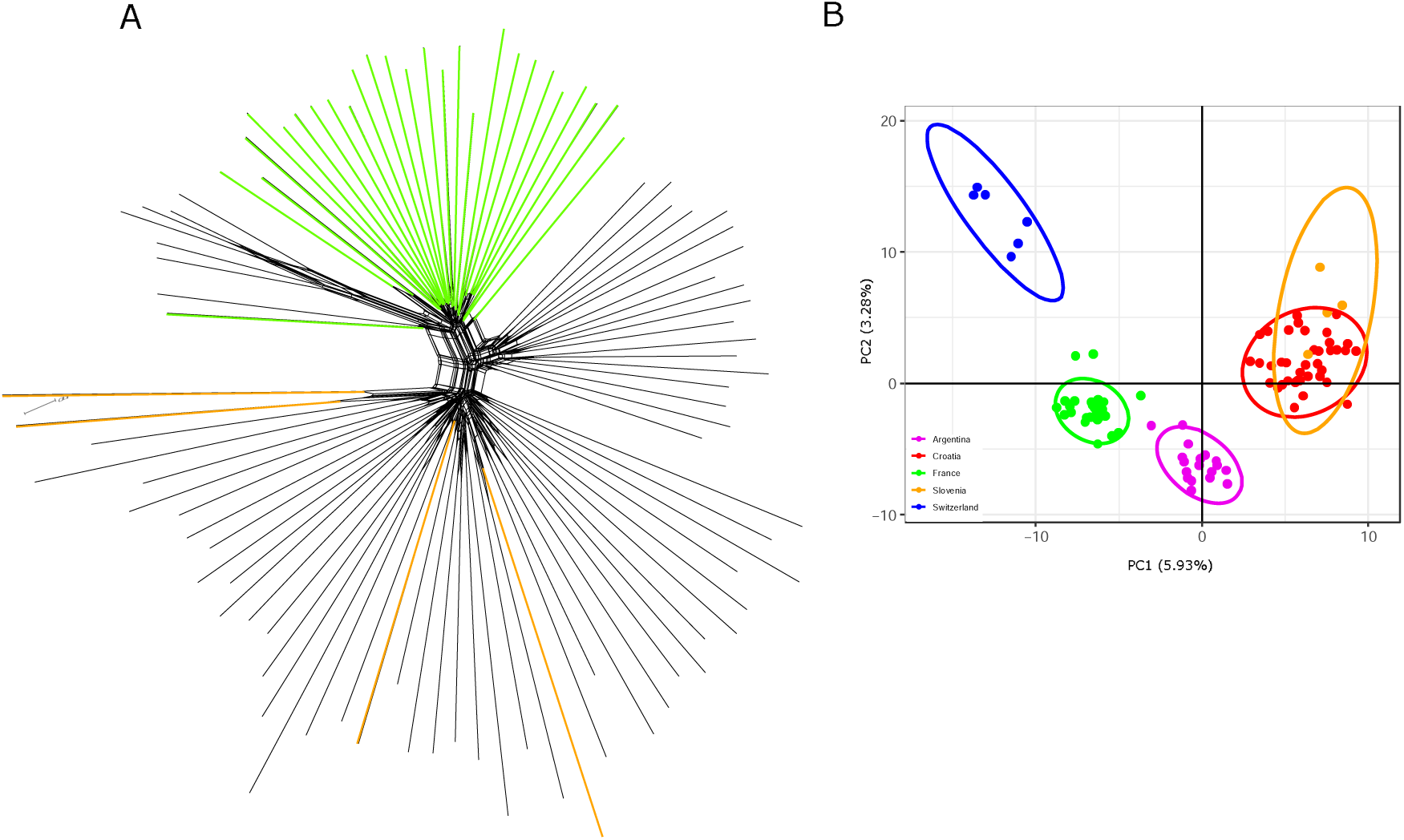
The European clade of *Colletotrichum graminicola.* (A) Neighbor-net network showing relationships between European isolates. (B) Principal-components analysis (PCA) with *a priori* geographical knowledge from sampling by country.

The relationship between all genotypes visualized in a neighbor phylogenetic network (Fig. S3) supported a clear worldwide subdivision into three lineages, with long branches separating lineages by geographic origin. We observed several loops (or reticulation) in the network indicating phylogenetic inconsistencies, where part of the focal sequence resembles that of a second sequence and one part of that sequence is more like a third sequence. Such a pattern is likely to be caused by recombination, as evidenced by the pairwise homoplasy index (PHI test) also calculated using SPLITSTREE which detected statistically significant evidence for recombination for the three lineages (*P*<0.001).

The consensus tree generated by DENSITREE, based on 1291 SNPs on chromosome 1, revealed many different topologies. This representation further corroborated the inferred population subdivision into three groups, with individuals from different lineages consistently separated by consensus branches (Fig. S5). We observed that isolates from different continents appear to have exchanged partial genomic sequences, indicating that part of genomes has a different phylogenetic origin, consistent with gene flow and recombination between diverged lineages (i.e., genetic introgression). Such events are illustrated by the diagonal lines of terminal branches in the DENSITREE cladogram.

### Gene flow analysis

Isolation-by-distance (IBD) analyses reveal a positive relationship between genetic and geographic distance. In total, 35.80% of the genetic diversification is explained by the geographical distance between sampling locations (Quadratic Regression F_2,40142_ = 11196.59, *P*<0.001, R^2^=0.3580) (Fig. 3). The isolation-by-distance plot shows that as geographic distance increases, the genetic differentiation between isolates becomes much more pronounced, but that it reaches a plateau. In other words, samples separated by a medium geographic distance (i.e., in the range of countries) are similarly diverged as those located on different continents. In addition, the Mantel test revealed a significant correlation between the genetic and geographic matrices (R^2^=0.3607, *P*<0.001). When we split the dissimilarity matrices into two groups according to geographical distance, we can see a clear pattern of isolation. Group 1 encompasses pairwise comparisons of samples separated by less than 3500 km (typical of isolates sampled on the same continent), and this shows the well-known isolation-by-distance relationship. This pattern reflects the natural gene flow of the pathogen. However, pairwise comparisons of samples separated more than 3500 km showed a very interesting pattern whereby the relationship reverses (Group 2). This negative correlation in isolation-by-distance likely reflects an artificial gene flow between distant populations caused by human-mediated transport. When analyzing these groups separately, we found a significantly positive IBD pattern at the inter-continental scale of <3500km (Regression: F_1,4843_=572.05, *P*<10^-6^, R^2^=10.6%), and a significantly negative IBD pattern between-continental scale of >3500km (F_1,8848_=3804.2, *P*<10^-6^, R^2^=30.1%).

**Figure 3.**
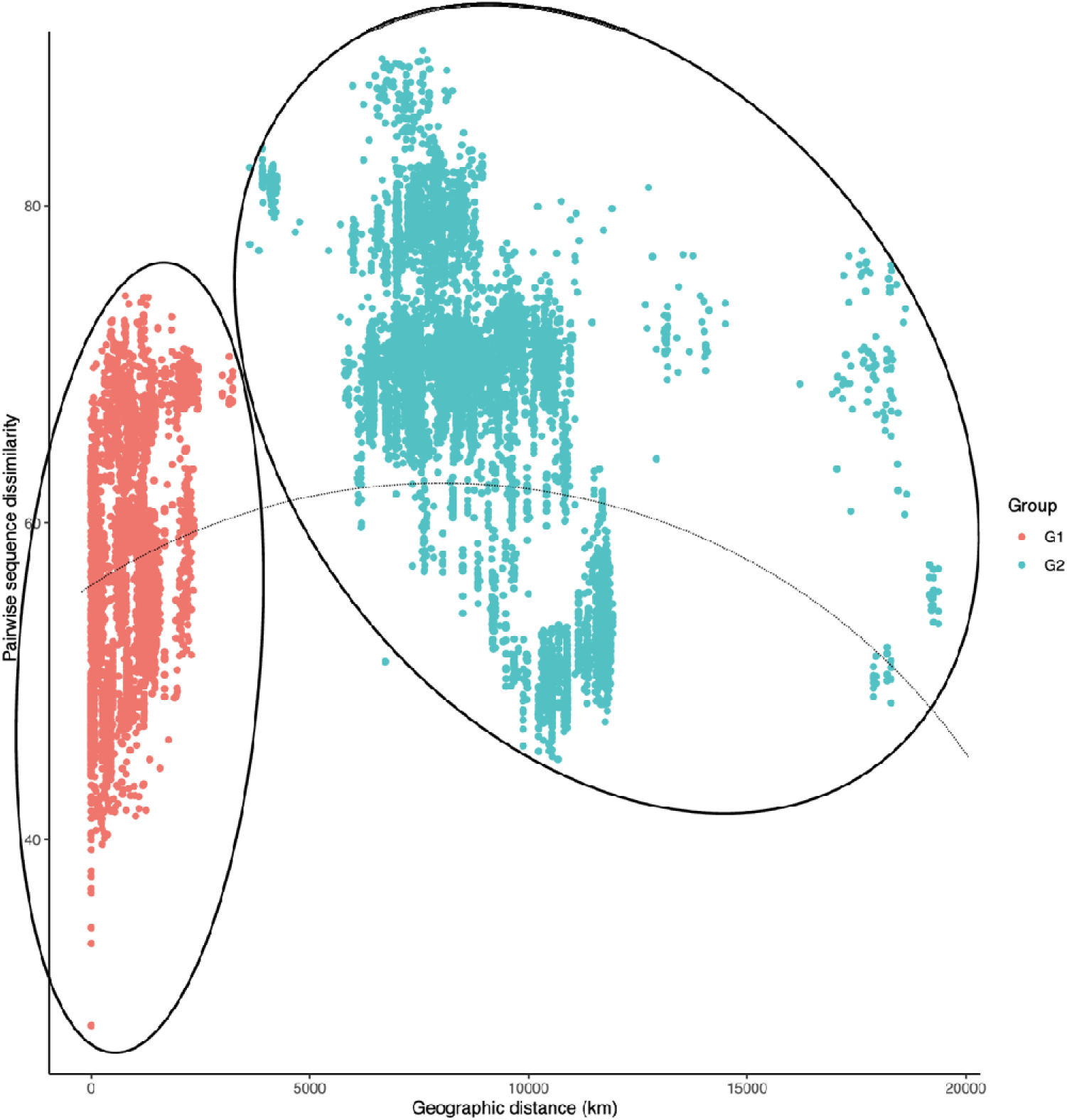
Isolation-by-distance plot showing the correlation between pairwise genetic dissimilarity and the geographical distance (in km). Quadratic regression F_2,40142_ = 11196.59, *P* < 0.001, R^2^ = 0.3580. Red-colored dots indicate Group 1 (geographical distance less than 3500 km, representing isolates sampled on the same continent) and the blue dots indicate Group 2 (geographical distance more than 3500 km, typical of samples from different continents).

### Recombinant analysis

Recombination in chromosome 1 was analyzed using RDP4, focussing on fifteen isolates representative of each lineage (North America: LAR318, NRRL13649, I-618151, A-52621-1, CA-CHAT-1; Brazilian: BR-85955-2, BR-98290-1, BR-85925-1, M5.001, JAB2; European: F-64330-2, ARG-X5196-1, P-7565-072-8, CR-34543-1, CBS11373). We detected 15 significant recombined blocks with large stretches of nucleotide similarity across lineages (Table S5).

Among the recombination events detected, we selected two (events 2 and 6) for more detailed analysis. We selected these events to illustrate the genomic signature of inter-continental genetic introgression, noting that the other 13 events show a comparable (albeit less clear) signature. Event 2 involves isolates BR-85955-2 (Brazilian lineage), P-7565-072-8 (European lineage), I-61851 (North American lineage), – henceforth referred to as triplet 1. In addition, event 6 consists of the isolates M5.001 (Brazilian lineage), CR-34543-1 (European lineage), A-52621-1 (North American lineage), henceforth to at triplet 2. Next, we visualized the recombinant blocks and dated their divergence using HYBRIDCHECK (Fig. 4). The visualization of nucleotide similarities between sequences through an RBG color triangle revealed a mosaic-like genome structure in these triplets (Ward and van Oosterhout 2016). Rather than a single recombination block as detected by RDP4, the blocks appear to be fragmented, possibly due to subsequent recombination and/or new mutations. The age estimates of those recombinant blocks varied widely among blocks (Table S6). We observed younger blocks dating back to 6,100 years before the present, and the older block sharing a common ancestor one dated to 100,000 years before the present.

**Figure 4.**
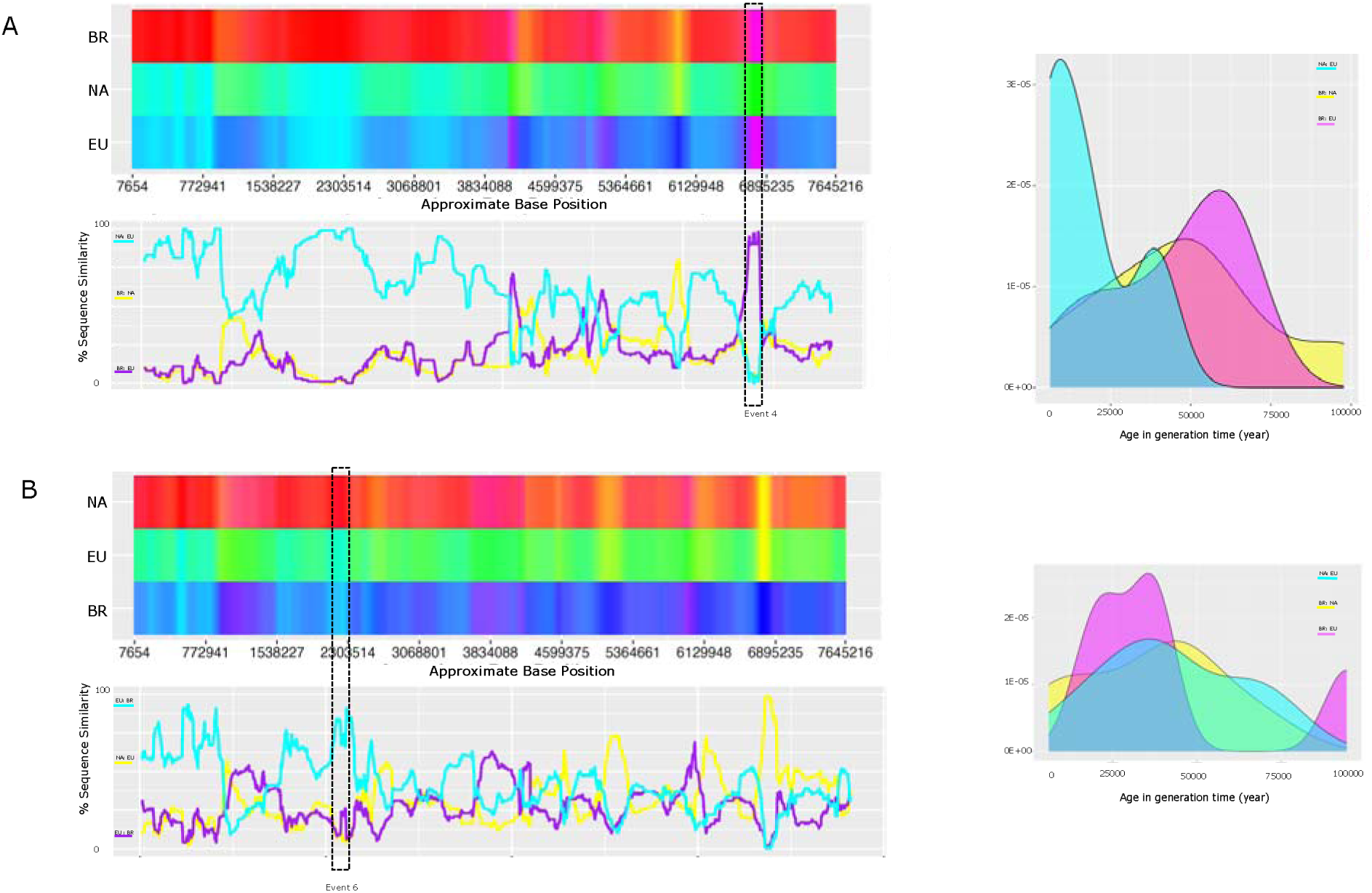
Recombination analysis of chromosome 1. (A) Left: Sequence similarity among the North American lineage (NA), European lineage (EU), and Brazilian lineage (BR), visualized through an RBG color triangle by HYBRIDCHECK (involving the isolates BR-85955-2, P-7565-072-8, and I-61851, respectively). Significant recombinant event detected by RPD4 software is enclosed in a dashed box. Right: Age distribution of recombinant blocks detected by HYBRIDCHECK. (B) Similar analyses of BR, EU, and NA lineages using the isolates M5.001, CR-34543-1, and A-52621-1, respectively.

To further investigate the variation of introgression age on the intra-continental scale, we performed an additional recombination analysis. Isolates from the same lineage were organized in three further triplets: triplet3 - M5.001:BR-98290-1:BR-85925-1 (Brazilian lineage), triplet 4 - F-64330-2:F-64330-7:F-64330-13B (European lineage), triplet 5 - NRRL13649:I-61851:CA-CHAT-1 (North America lineage) (Fig. S6). The recombinant blocks detected within each lineage were dated using HYBRIDCHECK and a Bayesian coalescent approach implemented in BEAST. The results indicated that the number of recombinant blocks and their age estimates differed significantly between lineages (Kruskal–Wallis test, H=33.5, d.f.=2, *P*<0.001). The age estimates provided by HYBRIDCHECK revealed that the blocks involving Brazilian and European isolates were younger, i.e., 25,000 and 30,000 years respectively (Fig. S7). In contrast, the North American isolates exhibited a more extensive number of blocks, with a much older signature of recombination, with some blocks dating nearly 100,000 years (Table S6 and Fig. S7).

To apply the Bayesian dating method, we selected three recombination blocks within each triplet identified by HYBRIDCHECK (for the blocks selected, refer to Table S6 - highlighted in yellow). These blocks were aligned against five isolates from each lineage to check whether they were conserved in the isolates (Table S7). Interestingly, these blocks were present in all isolates, displaying a pairwise identity greater than 99%. These sequences were used to estimate their age using BEAST. The dating results exhibited consistency with the HYBRIDCHECK algorithm dating, although there were differences in the order of magnitude of dates between the methods (Table S6 and Table S7). The age estimates for the North American (NA) blocks were more than seven times older than blocks from Brazil, ranging from 90,000 to 1,048,860 years (Table S7), a highly significant difference (Kruskal-Wallis test: H=32.37; d.f.=2; *P*<0.001). This suggests that the NA lineage has experienced a more ancient introgression than the other two lineages. We note that although the age of introgression may be overestimated, for the reasons explained in the Material and Methods, the relative difference in age estimates is noteworthy and likely to represent genuine differences in the history of introgression between the three geographic clusters.

### Unraveling population history

We analyzed the evolutionary history of the isolates assuming three genetic lineages North American (NA), Brazilian (BR), and European (EU) as independent populations. In total, we analyzed 22 demographic models (Fig. S8) using Approximate Bayesian Computation (ABC) based on random forest (RF), conducted in the Python package PYABCRANGER (Collin, et al. 2020). The demographic models tested were organized into four independent comparisons, to determine the most probable ancestral population (Fig. S9, and Fig. S10). Overall, our demographic modeling analysis strongly indicates a significant contribution of an unsampled population as the ancestral source shaping the complex evolutionary history of the lineage investigated. However, the North American population appears to have emerged first followed by the Brazilian and the European populations (Fig. 5 and Supplementary text S1). North America seems to have acted as an intermediate between the unknown source population and the rest of the world.

**Figure 5.**
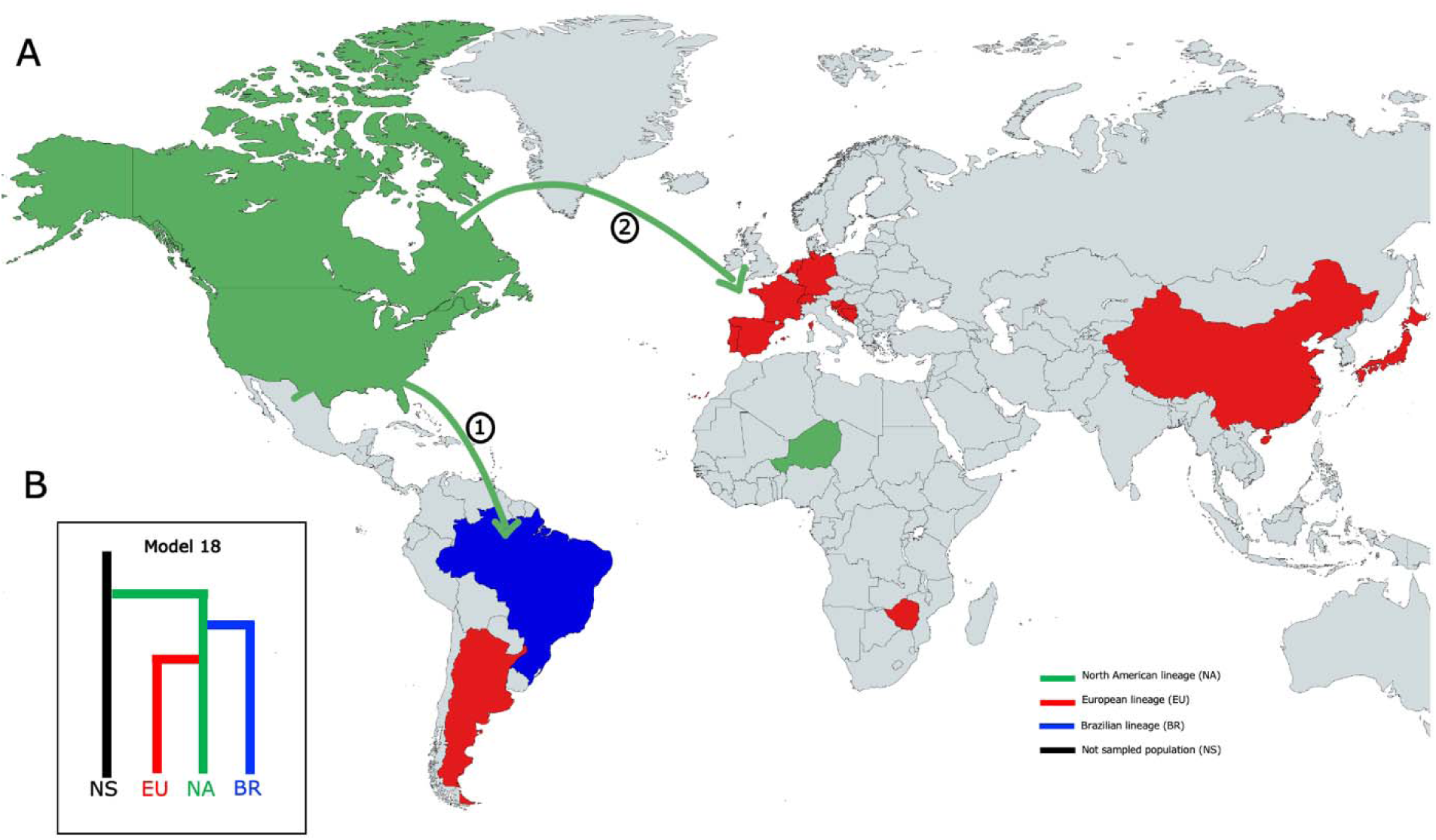
Worldwide sampling of *Colletotrichum graminicola*. (A) Map showing a likely colonization route of *Colletotrichum graminicola* reconstructed by Approximate Bayesian Computation (ABC) using 208 isolates from 17 countries. Different colors indicate distinct lineages and arrows indicate divergence events. **(1)** First divergence event from NA to BR; **(2)** Second divergence event from NA to EU. (B) Strongly supported demographic models voted based on random forest population (see Supplementary text S1 for details).

### Virulence analysis

Interestingly, we detected significant differences in virulence among the fifty-three isolates evaluated in an analysis of variance test (ANOVA, *P*<0.0001). Sidäk’s comparison-of-means test revealed that 16 isolates exhibited significantly higher virulence, while 14 isolates demonstrated significantly lower virulence compared to M1.001 (Fig. 1a and Fig. S11). The isolates CBS252.59, MAFF511343, and DMSZ-63127 were nonpathogenic to the maize cultivar evaluated.

The isolates were also classified according to genetic lineage (NA, EU, and BR) and sampling year. Differences in virulence between lineages were not statistically significant, based on a Kruskal-Wallis test, although greater variability was observed in the European group (Fig. S12a). We used as a control (or baseline) the strain M1.001, which was originally collected from North America in 1972 during the US outbreak and is highly virulent (Rogério et al. 2023; O’Connell et al. 2012), to classify the isolates into three categories (less, equal, or more virulent than isolate M1.001; see Table S10). When the isolates were cross-tabulated into these categories the percentages of strains in the different lineages were statistically significant (Fisher exact test, *P* < 0.05). All the isolates from the Brazilian lineage (BR) are less than or equally virulent than the strain M1.001, while 41.2% of the European lineage (EU) and 20.0% of the North American lineage (NA) are more virulent. These results indicate that there is a relationship between the genetic structure and virulence, particularly showing strains that are now more virulent than M1.001, especially in the European lineage. When the isolates were grouped by year (before 2012, between 2012 and 2016, and after 2016), a significant difference was observed between groups (Kruskal–Wallis test, H=9.02, d.f.=2, *P*=0.01). Isolates from 2012 to 2016 showed the highest virulence suggesting that within the three genetic groups, the relative level of virulence can change over time (Fig. S12b).

The Mantel test performed between pairwise genetic distance and pairwise similarity in virulence revealed no correlation between these variables (*R*=-0.006257, *P*=0.47053).

Interestingly, the isolates SW-8046-1 and SW-8046-2 exhibit (near) identical genotypes, yet they display contrasting virulence values (Fig. 1a and Fig. S11), suggesting that variations in virulence may be attributed to factors other than genetic diversity.

## Discussion

A global collection of *Colletotrichum graminicola* field isolates from 17 countries allowed a comprehensive investigation into the evolutionary history of this important maize pathogen. We generated and analyzed a broad data set with a higher resolution than our previous study (Rogério et al., 2023), supporting the three geographically near-isolated populations restricted, largely defined by continental origin. We identified an intriguing pattern of isolation-by-distance (IBD) consistent with two distinct gene flow patterns. A short-distance (<3500km) pattern of gene flow resulted in a typical IBD pattern time (Charlesworth 2009; Wright 1943), indicative of natural dispersal, possibly mediated by asexual spores. However, long-distance (>3500km) dispersal, showed a reverse pattern, i.e., a negative correlation between genetic and geographic distance. We interpret this as evidence of inter-continental human-mediated gene flow, possibly resulting from the trade and transport of contaminated seeds. Recombination analysis unveiled compelling evidence of genetic introgression between the three geographic groups, suggesting a long history of recombination, possibly through sexual reproduction. The North American lineage displayed the most ancient evidence of recombination. Demographic modeling supported these findings, indicating that North America is an intermediate between the source population and the global distribution, potentially originating from an unsampled population, hypothesized to be Mesoamerican. We propose that the global genetic structure of *C. graminicola* is shaped by geographic differentiation driven by long-distance migration and genetic recombination events occurring at distinct points in time.

Natural gene flow in fungi tends to be mediated through the transport of spores. The *Colletotrichum* genus typically produces asexual spores (conidia) in acervuli immersed in mucilaginous mass, disseminated through water splashes, accounting for short-range dispersal (Madden 1997). In contrast, sexual spores (ascospores) are associated with medium to long-distance dissemination, as they can be ejected upwards and dispersed by air currents, reaching longer distances (Taylor et al. 1999; Alaniz et al. 2019). Sexual reproduction in *C. graminicola* has the potential (even sporadically) to contribute to medium to long-distance gene flow (Rogério et al., 2023), given the prevailing assumption that asexual reproduction serves as the primary reproductive strategy for short-distance fungal dissemination (Drenth et al. 2019).

Human-mediated transport is a major driver of the long-distance dissemination of plant pathogens, acting as an artificial gene flow mechanism in scenarios where fungal spores may not naturally reach (Finlay 2002). The movement of contaminated seeds and infected plant material, driven by global human trade, can facilitate the migration of pathogens. This artificial gene flow may promote the movement of individuals and their alleles from one population to another, allowing genetic exchange between samples separated continentally, particularly among countries with strong trading ties. On a global scale, human-mediated migration is primarily responsible for the geographic differentiation of numerous populations of plant pathogens (Goss et al. 2014; Chakraborty 2013; Tabima et al. 2020; Kamvar et al. 2015b; Stukenbrock et al. 2006; McMullan et al. 2018; Linde et al. 2002; Zaffarano et al. 2006; Sotiropoulos et al. 2022; Barrès et al. 2008).

We detected migration between Argentina and Europe, where isolates from Argentina were grouped within the European lineage. This suggests some form of genetic exchange between these geographically distant regions. The role of contaminated seeds as sources of inoculum, promoting the movement of pathogens, has been well-established in this pathosystem (Rogério et al., 2023). In this mentioned study, we proposed the hypothesis that winter nurseries in South America, frequently employed in breeding programs, might serve as a potential source for the introduction of the pathogen into new regions. Furthermore, there seems to be a recurring pattern of maize germplasm movement between international breeding companies and experimental stations in Argentina. A case in point is the experimental station “La Josefina S.A”, in Mercedes, Argentina, which provided Winter Nursey Service from 1985 to 2014, collaborating with numerous (10 to 15 companies/institutes) maize breeding programs, including European (J. Murad 2023, personal communication). This station routinely handled maize seeds from international companies, working with germplasm from Europe, the USA, and Argentina. Consequently, it regularly received and returned seeds to the originating breeding programs, potentially contributing to the long-distance dissemination of the pathogen.

We identified signatures of genetic introgression at the genome-wide level between lineages, supported by statistically significant recombinant events. The examination of nucleotide similarities between sequences uncovered a mosaic-like genome structure, wherein different segments exhibit distinct genetic ancestries. The recombination analysis performed by RDP4 unveiled few, large recombinant blocks, contrasting with analyses using HYBRIDCHECK, which displayed numerous smaller, fragmented blocks. Through visualizing nucleotide similarities, we observed a pattern where smaller blocks appeared to be later fragmented by recurrent recombination. Multiple mutations may have accumulated after ancient recombination (i.e., before divergence), giving rise to the observed pattern of fragmented blocks. In essence, these small regions might be remnants of a common ancient introgression event, where subsequent recombination has fragmented the originally larger block.

It is important to note that subsequent recombination may also increase the genetic divergence between the donor and recipient sequence, making the introgressed regions appear more diverged and hence “older”. We are therefore cautious in our interpretation of the actual estimated divergence time dates. Nevertheless, we are confident in our interpretation that the introgression events observed in the North American lineage date further back in time than those in the other two lineages (i.e., of Europe and Brazil). Our recombination analysis detected events that took place as recently as 6100 years ago, as well as potentially very old events (>100,000 years ago). These more recent occurrences are likely indicative of genetic exchanges that have occurred in a relatively recent timeframe if not an ongoing process. Overall, our introgression investigation reveals a prolonged history of genetic exchanges.

Sexual reproduction has been observed under laboratory conditions, suggesting an absence of prezygotic reproductive barriers between genetically divergent strains (Rogério et al., 2023), as observed in other plant pathogenic ascomycetes (Le Gac and Giraud 2008). Although the sexual state of this species has not been documented in nature, our data suggest a significant contribution of genetic recombination, resulting from either sexual recombination or other non-meiotic recombination mechanisms, such as parasexual events via hyphal anastomosis, as previously described in other *Colletotrichum* species (Roca et al. 2003; Rosada et al. 2010; Souza-Paccola et al. 2003; Vaillancourt et al. 2000; Nordzieke 2022).

We propose that *C. graminicola* populations may have undergone intense recombination before genetic divergence, likely via sexual reproduction. Repeated sexual recombination cycles over time could have led to the development of a mosaic-like genome structure, as evidenced by our analyses. Ancient hybridization events have been observed in the wheat powdery mildew pathogen *Blumeria graminis* f. sp. *tritici*, characterized by admixture between divergent strains resulting in a mosaic genome. Chromosomal segments inherited from both parental sources exhibit fragmented segments, likely produced by backcrossing (Menardo et al. 2016; Sotiropoulos et al. 2022). A similar mosaic-like genome structure has been observed in the oomycete, *Albugo candida* (McMullan et al. 2015).

Regarding the evolutionary origin of these lineages, our demographic inference indicates that the North American lineage emerged before the Brazilian and European populations. Age estimates of recombinant blocks detected within lineages further support this finding. Dating the blocks involving isolates from North America revealed that this lineage is older compared to those blocks from other lineages. Moreover, the deeper branches in the phylogenetic tree and the higher level of genetic diversity are all consistent with the North American lineage predating the European and Brazilian lineages.

North America seems to act as an intermediate between the source population and the rest of the world, suggesting a potential colonization route of the fungus. Our results strongly indicate a significant contribution of an unsampled population as the ancestral source. Given the widely accepted hypothesis that maize domestication originated in Mexico (Sawers and Sanchez Leon 2011; Matsuoka et al. 2002), and following the principle that the center of origin of a pathogenic species usually corresponds to that of its host, that unsampled population might indeed represent a Mesoamerican population. However, our sampling did not include isolates from Mesoamerica, a region that could be linked to the ancestral population. To date, attempts to obtain isolates of *C. graminicola* from Mexico have been unsuccessful, and personal communications with researchers have revealed that it is not a disease of major importance in the region.

Recent studies indicate that maize underwent migration out of Mexico in two waves, at different time points (Wade 2023; Yang et al. 2023). Following the maize dispersal route, these findings suggest that two (or more) ancestral lineages may have contributed to the genetic diversity of this fungal species. After domestication, maize spread along the Pacific coast, initially reaching Central America (Panama), and then extending South America, particularly the Andean region of Peru, and potentially Brazil (Kistler et al. 2018). Maize then appears to have moved northward, reaching southwestern North America, before reaching Europe (Yang 2023). In Europe, maize was initally introduced into Spain from the Caribbean area in 1494 and then spread through the Mediterranean area, Africa and Asia. Additionaly, it was also introduced through the European Atlantic coast from North America (Revilla et al. 2022, 1998). However, due to the lack of comprehensive fungal sampling from South America, this region may have originated from the Mesoamerican *C. graminicola* lineage first, given the historical trajectory of the maize movement.

Establishing the origin of emerging infectious diseases is both challenging and important. Identifying the ancestral population reveals the standing genetic variation that could potentially introgress into the outbreak lineages. We encourage future studies to comprehensively survey across the Mesoamerica region, as well as South America, which would further advance our understanding of how this pathogen has adapted and spread to other parts of the world. A genomic comparison between the genetic variation in the source population and population worldwide would enable the examination of genes and genomic regions that experienced strong selection in the invasive strains. Such an analysis could shed light on the role of recombination and hybridization, which become more powerful when there is a greater presence of ancestral variation.

Finally, we observed great variability in virulence among the isolates tested, suggesting potential fungal adaptive evolution. A significant, albeit subtle, relationship was found between genetic groups and virulence levels, with the European group exhibiting increased virulence compared to the strain collected during the US outbreak. This level of virulence changed over time, raising concerns about potential new outbreaks of maize anthracnose, particularly in European cultivation. However, the absence of correlation between the pairwise genetic distance and virulence distance suggests that gene expression, epigenetic factors, or other factors not considered in this study may influence the variation in virulence. In addition, we observed that two pairs of isolates with nearly identical genotypes showed marked differences in virulence, emphasizing that adaptive evolution may be attributed to factors other than genetic diversity, such as environmental influences or epigenetic modifications. Further studies using these genotypes with contrasting virulence phenotypes may be used to elucidate the genetic basis of virulence in pathogens, contributing to a better understanding of plant diseases and the development of durable control strategies.

## Acknowledgments

The authors also would like to thank the Supercomputing and Bioinnovation Center (SCBI) of the University of Malaga (http://www.scbi.uma.es/site) and the Core Cluster of the Institut Français de Bioinformatique (IFB) (ANR-11-INBS-0013) for their provision of computational resources and technical support. We also wish to thank numerous colleagues who contributed to this work by providing samples of maize showing symptoms of anthracnose. We also thank Lucía Rodríguez Mónaco for her assistance during the virulence analysis.

## Data availability

The raw Illumina RAD-seq and WGS reads were submitted to GenBank and their accession numbers are listed in Table S2.

## Authors contributions

S.A.S., M.R.T. and F.R. conceived the study and designed the project; S.A.S., F.R. and S.G.-S. performed fungal isolation and DNA extraction; F.R., M.R.T., C.V.O., S.D.M., G.K.H., and J.L.V.-V. performed analyses; F.R., M.R.T., C.V.O., S.D.M. and S.A.S. wrote the manuscript; F.R., P.G.-R., S.B., S.G.-S., F.B.C.-F. and S.A.S. performed the pathogenic characterization; A.G.J., W.B., S.B.U., J.H., R.S., P.R., J.S.D., J.L.V. and I.B. collected the samples. All authors reviewed the manuscript.

## References

Alaniz, S., Cuozzo, V., Martínez, V., Stadnik, M. J., and Mondino, P. 2019. Ascospore infection and *Colletotrichum* species causing Glomerella leaf spot of apple in Uruguay. Plant Pathol J 35:100–111. 10.5423/PPJ.OA.07.2018.0145.

Anderson, E. 1953. INTROGRESSIVE HYBRIDIZATION. Biol Rev 28:280–307. 10.1111/j.1469-185X.1953.tb01379.x.

Ashu, E. E., Hagen, F., Chowdhary, A., Meis, J. F., and Xu, J. 2017. Global Population Genetic Analysis of *Aspergillus fumigatus*. mSphere 2:e00019–17. 10.1128/mSphere.00019-17.

Van der Auwera, G. A., Carneiro, M. O., Hartl, C., Poplin, R., del Angel, G., Levy-Moonshine, A., et al. 2013. From FastQ Data to High-Confidence Variant Calls: The Genome Analysis Toolkit Best Practices Pipeline. Curr Protoc Bioinformatics 43:11.10.1–11.10.33. 10.1002/0471250953.bi1110s43.

Barcelos, Q. L., Pinto, J. M. A., Vaillancourt, L. J., and Souza, E. A. 2014. Characterization of *Glomerella* strains recovered from anthracnose lesions on common bean plants in Brazil. PLoS One 9:e90910. 10.1371/journal.pone.0100438.

Barrès, B., Halkett, F., Dutech, C., Andrieux, A., Pinon, J., and Frey, P. 2008. Genetic structure of the poplar rust fungus *Melampsora larici-populina*: Evidence for isolation by distance in Europe and recent founder effects overseas. Infect Genet Evol 8:577–587. 10.1016/j.meegid.2008.04.005.

Barton, N. H. 2010. Mutation and the evolution of recombination. Phil Trans R. Soc B 365:1281–1294. 10.1098/rstb.2009.0320.

Becerra, S., Baroncelli, R., Boufleur, T. R., Sukno, S. A., and Thon, M. R. 2023. Chromosome-level analysis of the *Colletotrichum graminicola* genome reveals the unique characteristics of core and minichromosomes. Front Microbiol 14:1129319. 10.3389/fmicb.2023.1129319.

Beichman, A. C., Huerta-Sanchez, E., and Lohmueller, K. E. 2018. Using Genomic Data to Infer Historic Population Dynamics of Nonmodel Organisms. Annu. Rev Ecol Evol Syst 49:433–456. 10.1146/annurev-ecolsys-110617-062431.

Bergstrom, G. C., and Nicholson, R. L. 1999. The biology of corn anthracnose: Knowledge to exploit for improved management. Plant Dis 83:596–608. 10.1094/PDIS.1999.83.7.596.

Bhunjun, C. S., Phukhamsakda, C., Jayawardena, R. S., Jeewon, R., Promputtha, I., and Hyde, K. D. 2021. Investigating species boundaries in *Colletotrichum*. Fungal Divers 107:107–127. 10.1007/s13225-021-00471-z.

Bouckaert, R. R. 2010. DensiTree: Making sense of sets of phylogenetic trees. Bioinformatics 26:1372– 1373. 10.1093/bioinformatics/btq110.

Bouckaert, R. R., and Drummond, A. J. 2017. bModelTest: Bayesian phylogenetic site model averaging and model comparison. BMC Evol Biol 17:1–11. 10.1186/s12862-017-0890-6.

Bouckaert, R., Vaughan, T. G., Barido-Sottani, J., Duchêne, S., Fourment, M., Gavryushkina, A., et al. 2019. BEAST 2.5: An advanced software platform for Bayesian evolutionary analysis. PLoS Comput Biol 15:e1006650. 10.1371/journal.pcbi.1006650.

Byrnes, K. J., and Carroll, R. B. 1986. Fungi Causing Stalk Rot of Conventional-Tillage and No-Tillage Corn in Delaware. Plant Dis 70:238–239.

Carroll, D. 2013. Genetic Recombination. Second Edi. eds. Stanley Maloy and Kelly Hughes. San Diego: Academic Press. 10.1016/B978-0-12-374984-0.00627-6.

Casillas, S., and Barbadilla, A. 2017. Molecular population genetics. Genetics 205:1003– 1035. 10.1534/genetics.119.302623.

Chakraborty, S. 2013. Migrate or evolve: Options for plant pathogens under climate change. Glob Chang Biol 19:1985–2000.

Charlesworth, B. 2009. Effective population size and patterns of molecular evolution and variation. Nat Rev Genet 10:195–205. 10.1038/nrg2526.

Charlesworth, B. 2010. Molecular population genomics: A short history. Genet Res (Camb) 92:397–411.

Corsi, G. I., Tichkule, S., Sannella, A. R., Vatta, P., Asnicar, F., Segata, N., et al. 2022. Recent genetic exchanges and admixture shape the genome and population structure of the zoonotic pathogen *Cryptosporidium parvum*. Mol Ecol 1–13. 10.1111/mec.16556.

Cuevas-Fernández, F. B., Robledo-Briones, A. M., Baroncelli, R., Trkulja, V., Thon, M. R., Buhinicek, I., et al. 2019. First report of *Colletotrichum graminicola* Causing Maize Anthracnose in Bosnia and Herzegovina. Plant Dis 103:4–6. 10.1094/PDIS-06-19-1224-PDN.

Danecek, P., Auton, A., Abecasis, G., Albers, C. A., Banks, E., DePristo, M. A., et al. 2011. The variant call format and VCFtools. Bioinformatics 27:2156–2158. 10.1093/bioinformatics/btr330.

Dobzhansky, T. 1973. Nothing in biology makes sense except in the light of evolution. Am Biol Teach 35:125–129. 10.2307/4444260.

Drenth, A., McTaggart, A. R., and Wingfield, B. D. 2019. Fungal clones win the battle, but recombination wins the war. IMA Fungus 10:1–6. 10.1186/s43008-019-0020-8.

Everhart, S., Gambhir, N., and Stam, R. 2021. Population genomics of filamentous plant pathogens - A brief overview of research questions, approaches, and pitfalls. Phytopathology 111:12–22. 10.1094/PHYTO-11-20-0527-FI.

Felsenstein, J. 1974. The Evolutionary Advantage of Recombination. Genetics 78:737–756. http://www.jstor.org/stable/3276141.

Finlay, B. J. 2002. Global dispersal of free-living microbial eukaryote species. Science 296:1061–1063.

Frey, T. J., Weldekidan, T., Colbert, T., Wolters, P. J. C. C., and Hawk, J. A. 2011. Fitness evaluation of Rcg1, a locus that confers resistance to *Colletotrichum graminicola* (Ces.) G.W. Wils. Using near-isogenic maize hybrids. Crop Sci 51:1551–1563. 10.2135/cropsci2010.10.0613.

Frichot, E., Mathieu, F., Trouillon, T., Bouchard, G., and François, O. 2014. A short manual for sNMF: a program to estimate ancestry coefficients (command-line version). Genetics 196: 973–983. 10.1534/genetics.113.160572.

Giraud, T., Enjalbert, J., Fournier, E., Delmotte, F., and Dutech, C. 2008. Population genetics of fungal diseases of plants. Parasite 15:449–454. 10.1051/parasite/2008153449.

Gladieux, P., Byrnes, E. J., Aguileta, G., Fisher, M. C., Heitman, J., and Giraud, T. 2011. Epidemiology and Evolution of Fungal Pathogens in Plants and Animals. In Genetics and Evolution of Infectious Diseases, ed. Michel Tibayrenc. London: Elsevier, p. 59–132. 10.1016/B978-0-12-384890-1.00004-2.

Gompert, Z., and Buerkle, C. A. 2013. Analyses of genetic ancestry enable key insights for molecular ecology. Mol Ecol 22:5278–5294. 10.1111/mec.12488.

Goss, E. M., Tabima, J. F., Cooke, D. E. L., Restrepo, S., Frye, W. E., Forbes, G. A., et al. 2014. The Irish potato famine pathogen *Phytophthora infestans* originated in central Mexico rather than the Andes. Proc Natl Acad Sci USA 111:8791–8796. 10.1073/pnas.1401884111.

Jouet, A., McMullan, M., and Van Oosterhout, C. 2015. The effects of recombination, mutation and selection on the evolution of the *Rp1* resistance genes in grasses. Mol Ecol 24(12):3077–92. 10.1111/mec.13213.

Kamvar, Z. N., Larsen, M. M., Kanaskie, A. M., Hansen, E. M., and Grünwald, N. J. 2015. Spatial and Temporal Analysis of Populations of the Sudden Oak Death Pathogen in Oregon forests. Phytopathology 105:982–989. 10.1094/PHYTO-12-14-0350-FI.

Kamvar, Z. N., Brooks, J. C., and Grünwald, N. J. 2015b. Novel R tools for analysis of genome-wide population genetic data with emphasis on clonality. Front Genet 6:208. 10.3389/fgene.2015.00208.

Kasuga, T., White, T. J. and Taylor, J. W. 2002. Estimation of nucleotide substitution rates in eurotiomycete fungi. Mol Biol Evol, 19, 2318–2324. 10.1093/oxfordjournals.molbev.a004056

Kistler, L., Yoshi Maezumi, S., De Souza, J. G., Przelomska, N. A. S., Costa, F. M., Smith, O., et al. 2018. Multiproxy evidence highlights a complex evolutionary legacy of maize in South America. Science 362:1309–1313. 10.1126/science.aav0207.

Koskella, B., Giraud, T., and Hood, M. E. 2006. Pathogen Relatedness Affects the Prevalence of Within-Host Competition. American Naturalist 168:121–126. 10.1086/505770.

Letunic, I., and Bork, P. 2007. Interactive Tree Of Life (iTOL): An online tool for phylogenetic tree display and annotation. Bioinformatics 23:127–128. 10.1093/bioinformatics/btl529.

Li, H., and Durbin, R. 2009. Fast and accurate short read alignment with Burrows-Wheeler transform. Bioinformatics 25:1754–1760. 10.1093/bioinformatics/btp324.

Li, H., Handsaker, B., Wysoker, A., Fennell, T., Ruan, J., Homer, N., et al. 2009. The Sequence Alignment/Map format and SAMtools. Bioinformatics 25:2078–2079. 10.1093/bioinformatics/btp352.

Linde, C. C., Zhan, J., and Mcdonald, B. A. 2002. Population Structure of *Mycosphaerella graminicola*: From Lesions to Continents. Phytopathology 92:946–955. 10.1094/PHYTO.2002.92.9.946.

Lipps, P. E. 1983. Survival of *Colletotrichum graminicola* in infested corn residues in Ohio. Plant Dis 67:102–104. https://10.0.4.70/PD-67-102.

López-Villavicencio, M., Jonot, O., Coantic, A., Hood, M. E., Enjalbert, J., and Giraud, T. 2007. Multiple Infections by the Anther Smut Pathogen Are Frequent and Involve Related Strains. PLoS Pathog 3:1710–1715. 10.1371/journal.ppat.0030176.

Martin, D. P., Varsani, A., Roumagnac, P., Botha, G., Maslamoney, S., Schwab, T., et al. 2021. RDP5: A computer program for analyzing recombination in, and removing signals of recombination from, nucleotide sequence datasets. Virus Evol 7:veaa087. 10.1093/ve/veaa087.

Matsuoka, Y., Vigouroux, Y., Goodman, M. M., Sanchez, J. G., Buckler, E., and Doebley, J. 2002. A single domestication for maize shown by multilocus microsatellite genotyping. Proc Natl Acad Sci USA 99:6080–6084. 10.1073/pnas.052125199.

McDermott, J. M., and McDonald, B. A. 1993. Gene Flow in Plant Pathosystems. Annu Rev Phytopathol 31:353–373. 10.1146/annurev.py.31.090193.002033.

McDonald, B. A., and Linde, C. 2002. PATHOGEN POPULATION GENETICS, EVOLUTIONARY POTENTIAL, AND DURABLE RESISTANCE. Annu Rev Phytopathol 40:349–379. 10.1146/annurev.phyto.40.120501.101443.

McMullan, M., Gardiner, A., Bailey, K., Kemen, E., Ward, B. J., Cevik, V., et al. 2015. Evidence for suppression of immunity as a driver for genomic introgressions and host range expansion in races of *Albugo candida*, a generalist parasite. Elife 4:e04550. 10.7554/eLife.04550.

McMullan, M., Rafiqi, M., Kaithakottil, G., Clavijo, B. J., Bilham, L., Orton, E., et al. 2018. The ash dieback invasion of Europe was founded by two genetically divergent individuals. Nat Ecol Evol 2:1000–1008. 10.1038/s41559-018-0548-9.

Menardo, F., Praz, C. R., Wyder, S., Ben-David, R., Bourras, S., Matsumae, H., et al. 2016. Hybridization of powdery mildew strains gives rise to pathogens on novel agricultural crop species. Nat Genet 48:201–205. 10.1038/ng.3485.

Meng, J. W., Zhu, W., He, M. H., Wu, E. J., Duan, G. H., Xie, Y. K., et al. 2015. Population genetic analysis reveals cryptic sex in the phytopathogenic fungus *Alternaria alternata*. Sci Rep 5:1–10. 10.1038/srep18250.

Milgroom, M. G. 2015. Population Biology of Plant Pathogens: Genetics, Ecology, and Evolution. St. Paul, Minnesota.

Nordzieke, D. E. 2022. Hyphal Fusions Enable Efficient Nutrient Distribution in *Colletotrichum graminicola* Conidiation and Symptom Development on Maize. Microorganisms 10:1146. 10.3390/microorganisms10061146.

Nordzieke, D. E., Sanken, A., Antelo, L., Raschke, A., Deising, H. B., and Pöggeler, S. 2019. Specialized infection strategies of falcate and oval conidia of *Colletotrichum graminicola*. Fungal Genet and Biol 133:103276. 10.1016/j.fgb.2019.103276.

O’Connell, R. J., Thon, M. R., Hacquard, S., Amyotte, S. G., Kleemann, J., Torres, M. F., et al. 2012. Lifestyle transitions in plant pathogenic *Colletotrichum* fungi deciphered by genome and transcriptome analyses. Nat Genet 44:1060–1065. 10.1038/ng.2372.

Van Oosterhout, C. 2021. Mitigating the threat of emerging infectious diseases; a coevolutionary perspective. Virulence 12:1288–1295. 10.1080/21505594.2021.1920741.

Perfect, S. E., Hughes, H. B., O’Connell, R. J., and Green, J. R. 1999. *Colletotrichum*: A model genus for studies on pathology and fungal-plant interactions. Fungal Genet Biol 27:186–198. 10.1006/fgbi.1999.1143.

Revilla, P., Alves, M.L., Andelković, V., Balconi, C., Dinis, I., Mendes-moreira, P., et al. 2022. Traditional Foods From Maize (*Zea mays* L.) in Europe. Front nutr 8: 683399. 10.3389/fnut.2021.683399.

Revilla, P., Soengas, P., Malvar, R. A., Cartea, M. E., Ordás, A. 1998. ISOZYME VARIATION AND HISTORICAL RELATIONSHIPS AMONG THE MAIZE RACES OF SPAIN. Maydica, 43:175–182. https://digital.csic.es/handle/10261/42758.

Roca, M. G., Davide, L. C., Mendes-Costa, M. C., and Wheals, A. 2003. Conidial anastomosis tubes in *Colletotrichum*. Fungal Genet Biol 40:138–145. 10.1016/S1087-1845(03)00088-4.

Roca, M. G., Read, N. D., and Wheals, A. E. 2005. Conidial anastomosis tubes in filamentous fungi. FEMS Microbiol Lett 249:191–198. 10.1016/j.femsle.2005.06.048.

Rochette, N. C., Rivera-Colón, A. G., and Catchen, J. M. 2019. Stacks 2: Analytical methods for paired-end sequencing improve RADseq-based population genomics. Mol Ecol 28:4737–4754. 10.1111/mec.15253.

Rogério, F., Baroncelli, R., Cuevas-Fernandez, F. B., Becerra, S., Crouch, J., Bettiol, W., et al. 2023. Population genomics provide insights into the global genetic structure of *Colletotrichum graminicola*, the causal agent of maize anthracnose. mBio 14:e02878–22. 10.1128/mbio.02878-22.

Rogério, F., Van Oosterhout, C., Ciampi-Guillardi, M., Correr, F. H., Hosaka, G. H., Cros-Arteil, S., et al. 2022. Means, motive, and opportunity for biological invasions: genetic introgression in a fungal pathogen. Mol Ecol 32:2428–2442. 10.1111/mec.16366.

Rosada, L. J., Franco, C. C. S., Sant’anna, J. R., Kaneshima, E. N., Gonçalves-Vidigal, M. C., and Castro-Prado, M. A. A. 2010. Parasexuality in Race 65 *Colletotrichum lindemuthianum* Isolates. J Eukaryot 57:383–384. 10.1111/j.1550-7408.2010.00486.x.

Rosenzweig, B. K., Pease, J. B., Besansky, N. J., and Hahn, M. W. 2016. Powerful methods for detecting introgressed regions from population genomic data. Mol Ecol 25:2387–2397. 10.1111/mec.13610.

Sawers, R. J. H., and Sanchez Leon, N. L. 2011. Origins of maize: A further paradox resolved. Front Genet 2. 10.3389/fgene.2011.00053.

Siol, M., Coudoux, T., Ravel, S., and De Mita, S. 2022. EggLib 3: A python package for population genetics and genomics. Mol Ecol Resour 22:3176–3187. 10.1111/1755-0998.13672.

Sotiropoulos, A. G., Arango-Isaza, E., Ban, T., Barbieri, C., Bourras, S., Cowger, C., et al. 2022. Global genomic analyses of wheat powdery mildew reveal association of pathogen spread with historical human migration and trade. Nat Commun 13:1–14. 10.1038/s41467-022-31975-0.

Souza, E. A., Camargo Junior, O. A., and Casela, C. L. 2010. Sexual recombination in *Colletotrichum lindemuthianum* occurs on a fine scale. Genet. Mol. Res. 9:1759–1769. https://10.4238/vol9-3gmr863.

Souza-Paccola, E. A., Fávaro, L. C. L., Casela, C. R., and Paccola-Meirelles, L. D. 2003. Genetic Recombination in *Colletotrichum sublineolum*. J Phytopathol 151:329–334. 10.1046/j.1439-0434.2003.00727.x.

Steinrücken, M., Kamm, J., Spence, J. P., and Song, Y. S. 2019. Inference of complex population histories using whole-genome sequences from multiple populations. Proc Natl Acad Sci USA 116:17115–17120. 10.1073/pnas.1905060116.

Stukenbrock, E. H. 2016. The Role of Hybridization in the Evolution and Emergence of New Fungal Plant Pathogens. Phytopathology 106:104–112. 10.1094/PHYTO-08-15-0184-RVW.

Stukenbrock, E. H., Banke, S., and McDonald, B. A. 2006. Global migration patterns in the fungal wheat pathogen *Phaeosphaeria nodorum*. Mol Ecol 15:2895–2904. 10.1111/j.1365-294X.2006.02986.x.

Stukenbrock, E. H., and Bataillon, T. 2012. A Population Genomics Perspective on the Emergence and Adaptation of New Plant Pathogens in Agro-Ecosystems. PLoS Pathog 8:1–4. 10.1371/journal.ppat.1002893.

Stukenbrock, E. H., and Dutheil, J. Y. 2018. Fine-Scale Recombination Maps of Fungal Plant Pathogens Reveal Dynamic Recombination Landscapes and Intragenic Hotspots. Genetics 208:1209–1229. 10.1534/genetics.117.300502.

Sukno, S. A., García, V. M., Shaw, B. D., and Thon, M. R. 2008. Root infection and systemic colonization of maize by *Colletotrichum graminicola*. Appl Environ Microbiol 74:823–832. 10.1128/AEM.01165-07.

Tabima, J. F., Søndreli, K. L., Keriö, S., Feau, N., Sakalidis, M. L., Hamelin, R. C., et al. 2020. Population Genomic Analyses Reveal Connectivity via Human-Mediated Transport across Populus Plantations in North America and an Undescribed Subpopulation of *Sphaerulina musiva*. Mol Plant-Microbe Interact 33:189–199. 10.1094/MPMI-05-19-0131-R.

Talhinhas, P., and Baroncelli, R. 2021. *Colletotrichum* species and complexes: geographic distribution, host range and conservation status. Fungal Divers 110:109–198. 10.1007/s13225-021-00491-9.

Talhinhas, P., and Baroncelli, R. 2023. Hosts of *Colletotrichum*. Mycosphere 14:158–261. 10.5943/mycosphere/14/si2/4.

Taylor, J., Jacobson, D., and Fisher, M. 1999. THE EVOLUTION OF ASEXUAL FUNGI: Reproduction, Specification and Classification. Annu Rev Phytopathol 37:197–246. 10.1146/annurev.phyto.37.1.197.

Taylor, J. W., Hann-Soden, C., Branco, S., Sylvain, I., and Ellison, C. E. 2015. Clonal reproduction in fungi. Proc Natl Acad Sci USA 112:8901–8908. 10.1073/pnas.1503159112.

Vaillancourt, L., Wang, J., Hanau, R., Rollins, J., and Du, M. 2000. Genetic Analysis of Cross Fertility between Two Self-Sterile Strains of *Glomerella graminicola*. Mycologia 92:430–435. 10.2307/3761501.

Vargas, W. A., Sanz Martín, J. M., Rech, G. E., Rivera, L. P., Benito, E. P., Díaz-Mínguez, J. M., et al. 2012. Plant Defense Mechanisms aAe Activated during Biotrophic and Necrotrophic Development of *Colletotricum graminicola* in Maize. Plant Physiol 158:1342–1358. 10.1104/pp.111.190397.

de Vries, S., Stukenbrock, E. H., and Rose, L. E. 2020. Rapid evolution in plant–microbe interactions – an evolutionary genomics perspective. New Phytologist 226:1256–1262. 10.1111/nph.16458.

Wade, L. 2023. Maize has an unexpected wild ancestor. Science 382:983–984. https://10.0.4.102/science.adn2113.

Ward, B. J., and van Oosterhout, C. 2016. HYBRIDCHECK: software for the rapid detection, visualization and dating of recombinant regions in genome sequence data. Mol Ecol Resour 16:534–539. 10.1111/1755-0998.12469.

Wright, S. 1943. Isolation by distance. Genetics 28:114–138. 10.1093/genetics/28.2.114.

Yang, N., Wang, Y., Liu, X., Jin, M., Vallebueno-Estrada, M., Calfee, E., et al. 2023. Two teosintes made modern maize. Science 382:eadg8940. 10.1126/science.adg8940.

Zaffarano, P. L., McDonald, B. A., Zala, M., and Linde, C. C. 2006. Global hierarchical gene diversity analysis suggests the fertile crescent is not the center of origin of the barley scald pathogen *Rhynchosporium secalis*. Phytopathology 96:941–950. 10.1094/PHYTO-96-0941.

Zhan, J., and McDonald, B. A. 2013. Experimental Measures of Pathogen Competition and Relative Fitness. Annu Rev Phytopathol 51:131–153. 10.1146/annurev-phyto-082712-102302.

Zhan, J., Thrall, P. H., Papaïx, J., Xie, L., and Burdon, J. J. 2015. Playing on a Pathogen’s Weakness: Using Evolution to Guide Sustainable Plant Disease Control Strategies. Annu Rev Phytopathol 53:19–43. 10.1146/annurev-phyto-080614-120040.

