## Supplementary material for "Long-distance gene flow and recombination shape the evolutionary history of a maize pathogen": Suplementary text S1

**Supplementary text S1**

We inferred the evolutionary history of *Colletotrichum graminicola* genetic lineages using Approximate Bayesian Computation (ABC) based on random forest (RF), conducted in the Python package Pyabcranger (Collin, et al. 2020). We based our analysis on three genetic lineages (North American, Brazilian, and European) treated independent populations, following a previous study by Rogério et al. (2023). Five samples from Asia and Africa (JP-244463, CH-TZ-3, CBS113173, MAFF511343, LARS318) were removed from the observed dataset due to their external position on the neighbor network. As an outgroup, we used the species *Colletotrichum navitas* (strain CBS 125086; accession number SAMN05660908) mapped against genome reference and included in our dataset. Hence, the total of samples in this analysis equals 208 samples. We included in the analysis biallelic sites with a minor allele frequency threshold 5%, excluding sites with more than 50% of missing data in any population, and applied a thinning factor to generate a reference dataset of 745 SNPs spreading the whole genome.

We used the Python package egglib to perform coalescent simulations and to compute the summary statistics, for both observed and simulated datasets. In total 39 statistics were analyzed, which were calculated for the whole dataset, for individual populations, and/or for pairwise comparisons (Table S8). To validate the models employed, we performed a linear discriminant analysis (LDA) using the sklearn.discriminant_analysis function from the Python package scikit-learn (Pedregosa et al. 2011) on the summary statistics of simulated datasets and compare these against the summary statistics of the observed dataset (Fig. S9). In ABC analysis, Pyabcranger was used to perform model choice and parameter estimation. To identify the most probable model, ABC-RF uses a classification vote system after bagging (i.e., aggregating bootstrap results) of the simulated outputs. The model with the highest number of votes is deemed to be the most probable model. Posterior distribution of parameter estimates was estimated using 1,000,000 simulations. The expectation value, median, standard deviation and two quantiles were reported, and the distribution itself was obtained using kernel density evaluation (KDE) with scipty. The model choice was performed using subsets of 100,000 to 500,000 simulations per model.

We modeled several scenarios of divergence including asymmetrical migration and bottleneck events. Models with and without an unknown population (i.e., ‘ghost’ population) as an ancestral population were tested. In our models, every divergence event was immediately followed by a bottleneck, otherwise keeping constant population size for all populations. We compared 22 demographic scenarios assuming either stepping-stone and single-source models (Fig. S8; Table S9). Prior distributions of parameters were modeled as truncated normal distributions for each parameter, regardless of the model (see Table S9 for details).

First, we compared 13 models (model 1 to 13 - Additional file 1: Fig. S8) excluding an unsampled population to determine whether any of our populations could be identified as ancestral. Model 6 (NA ancestral, from which BR emerges first, followed by EU), consistently received the most votes in the random forest (130 votes, Posterior probability = 0.47) over the range of numbers of included simulations (Additional file 1: Fig. S10). Second, we investigated the potential ancestral role of an unknown ancestral population (NS) by comparing 9 models (models 14 to 22), including an unsampled population. Accounting for all possibilities being untractable, we restricted the analysis to models extending models best supported by the first analysis. For example, model 18 is equivalent to model 6 but NA ultimately emerges from NS while model 20 is an equivalent where BR and EU emerge from NS instead of NA. In this comparison, model 20 was the voted best (120 votes, posterior probability = 0.48), with a minimal gap with model 18 and then model 19 (where BR emerges from NS, then NA from BR, then EU from BR). Next, we conducted a comparison involving a subset of models, including best-ranking models from the first comparison (models 6,7,8,9 and 11) and models with an unsampled population (except model 17 in which population splits are simultaneous and which received the lowest number of votes in the second analysis). In this case, model 18 has the best number of votes, slightly better than model 19 (111 votes, posterior probability = 0.53). Finally, we compared two groups of models among the models used in the previous analysis: one consisting of models with an unknown population as ancestral (8 models) and the other one without (5 models). The group with an unknown population emerged as the clear preference, receiving 419 votes, and a posterior probability = 0.87.
