## Supplementary material for "Long-distance gene flow and recombination shape the evolutionary history of a maize pathogen": Suplementary figures S1-S12

Table of contents:


| | |
|---|---|
| Figure S1 | Page 2 |
| Figure S2 | Page 3 |
| Figure S3 | Page 4 |
| Figure S4 | Page 5 |
| Figure S5 | Page 6 |
| Figure S6 | Page 7 |
| Figure S7 | Page 8 |
| Figure S8 | Page 9 |
| Figure S9 | Page 10 |
| Figure S10 | Page 11 |
| Figure S11 | Page 12 |
| Figure S12 | Page 13 |

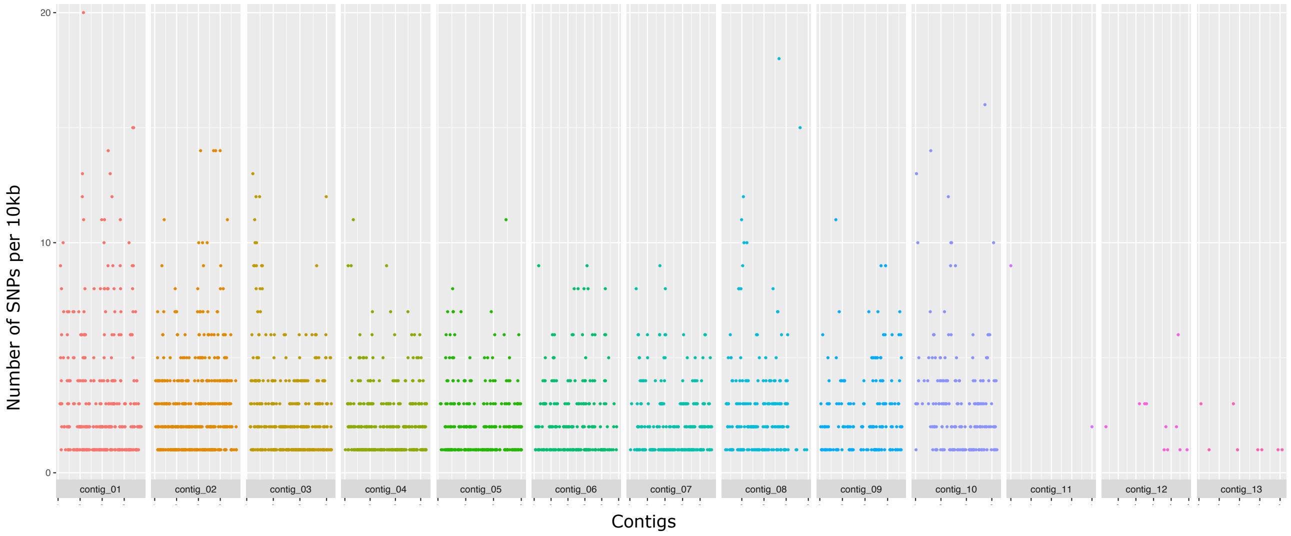


**Figure S1.** Manhattan plot of the distribution of the SNPs along the chromosomes. Each point represents the position of a SNP on the x-axis. SNP density was calculated per 10kb across chromosomes using vcftools software (Danecek et al., 2011).

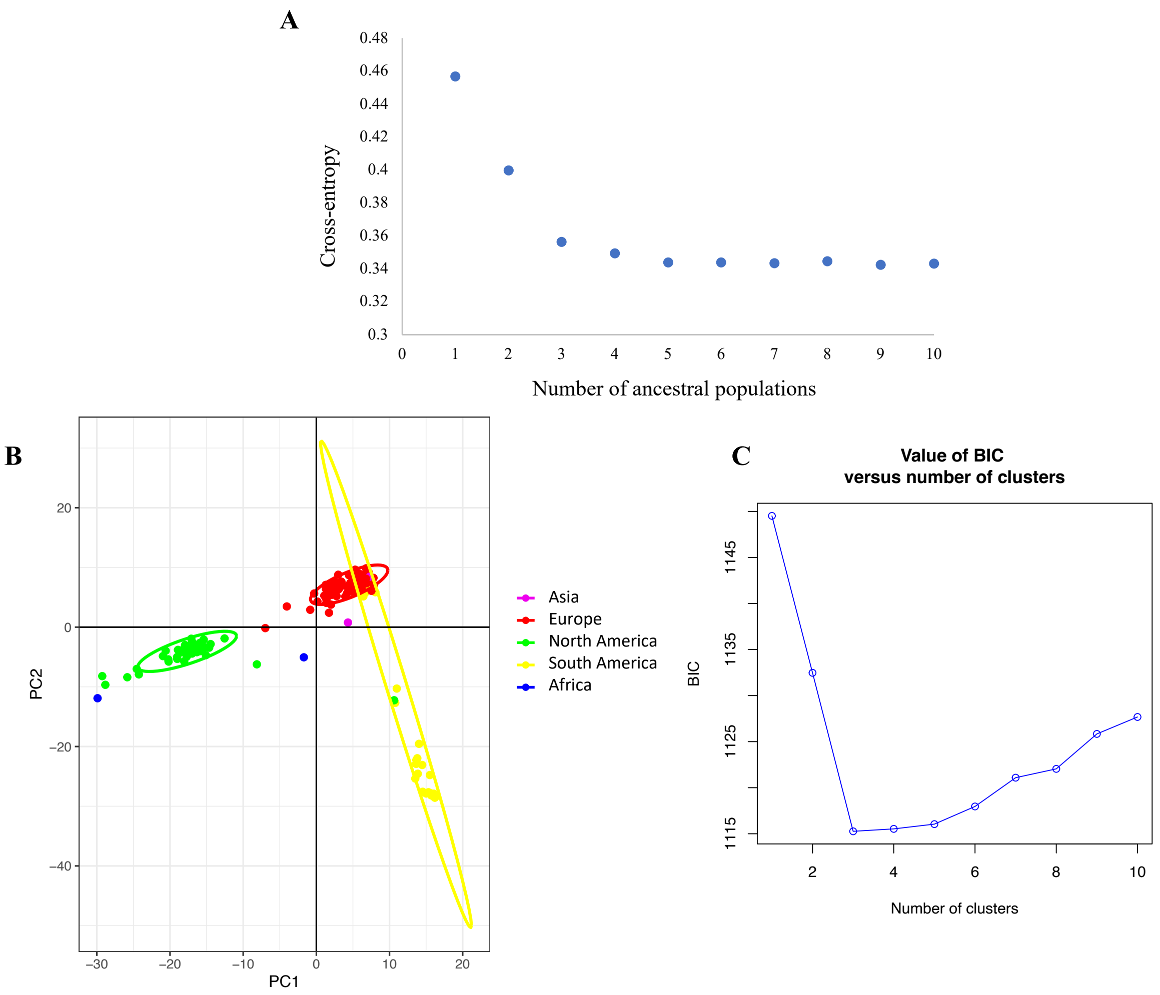


**Figure S2.** Population subdivision of *Colletotrichum graminicola.* (A) Cross-entropy as a function of the number of clusters *K* modeled in snmf analysis of population subdivision. (B) Principal-component analysis (PCA) with a priori geographical knowledge from sampling (by continent). (C) Bayesian information criteria (BIC) indicating the most probable number of genetic groups.

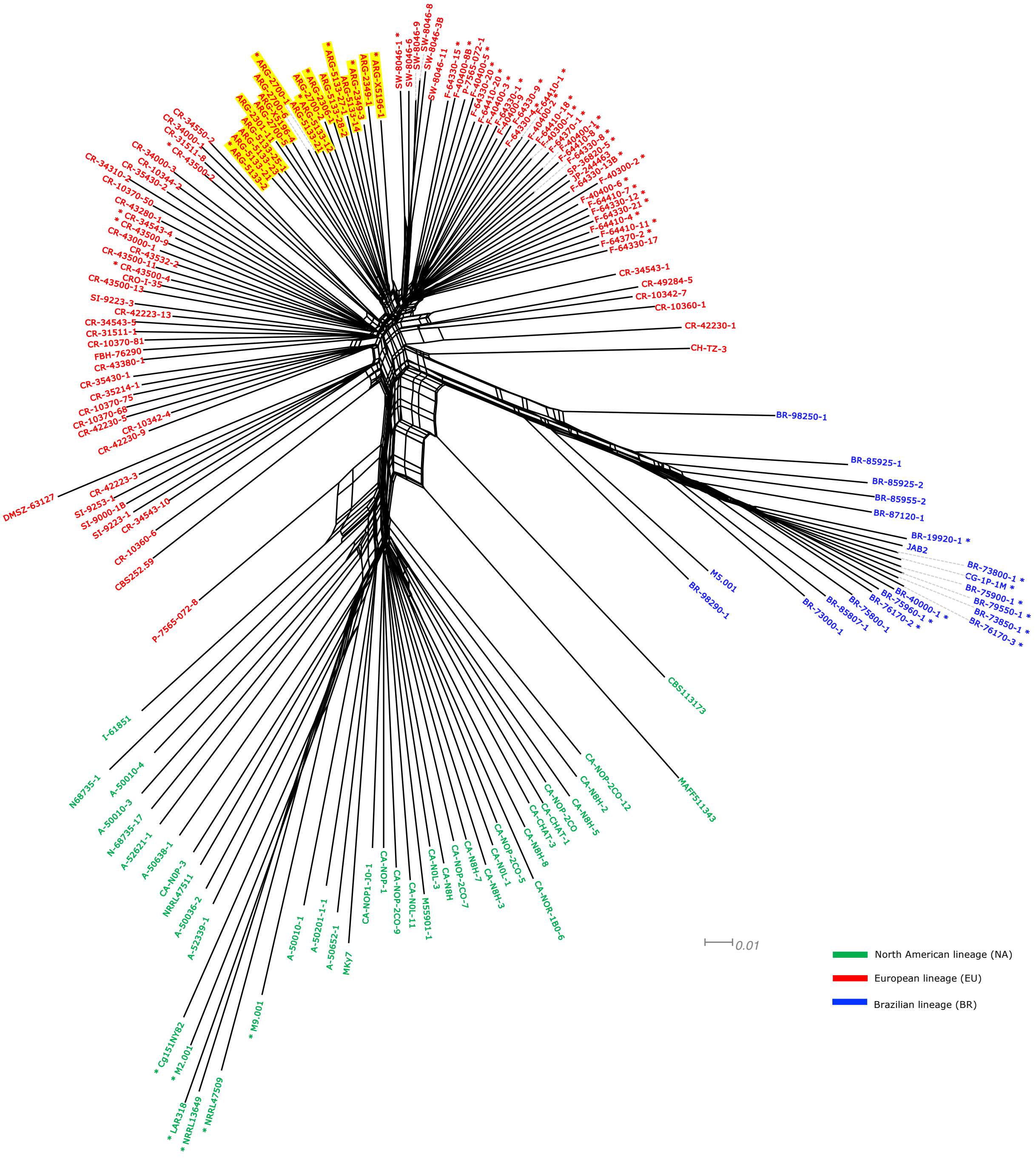


**Figure S3.** Neighbor-net network showing relationships between isolates of *Colletotrichum graminicola* identified based on the clone-correct dataset. highlights indicate “migrant”, i.e., isolates that cluster within the group of isolates from another geographic location. Asteristics indicate pure isolates ($q$ > 0.99).

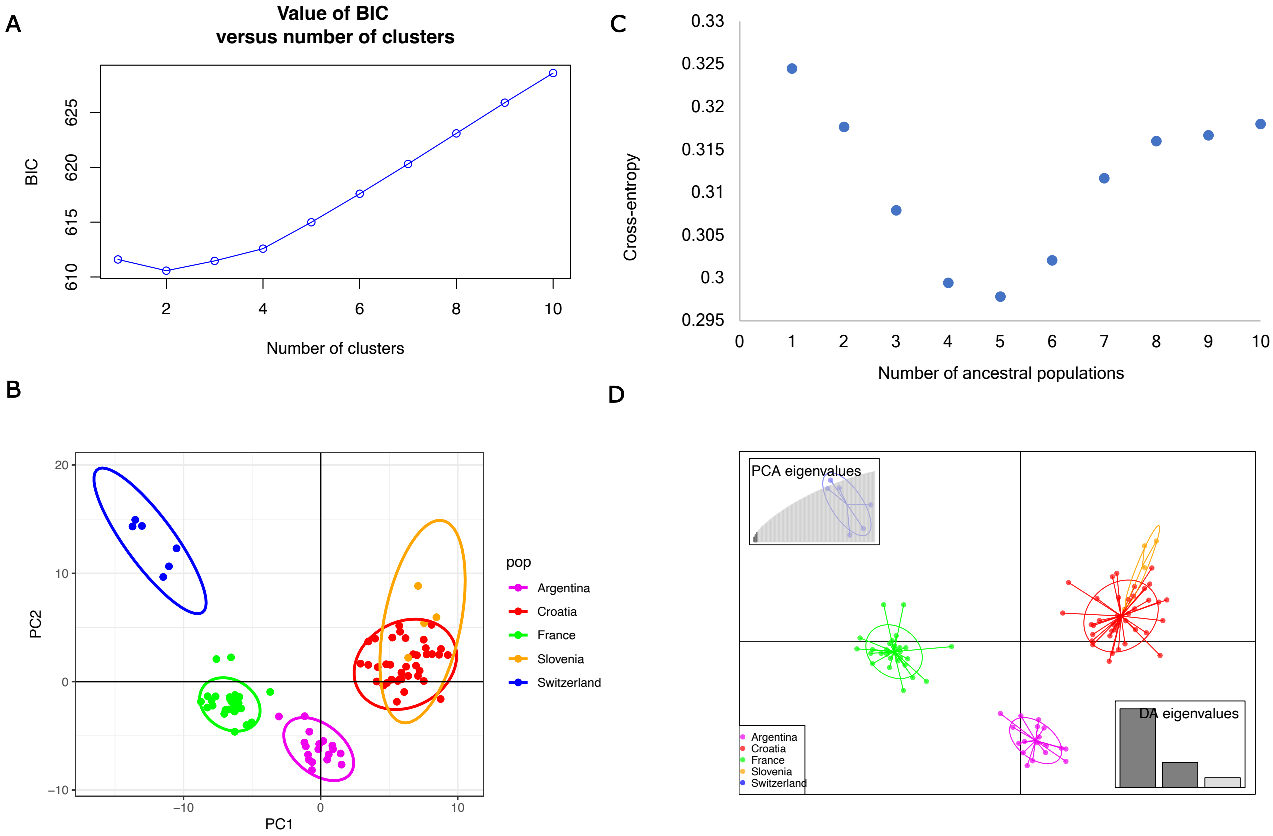


**Figure S4.** Population subdivision on European lineage of *Colletotrichum graminicola.* (A) Cross-entropy as a function of the number of clusters *K* modeled in snmf analysis of population subdivision. (B) Principal-component analysis (PCA) with a priori geographical knowledge from sampling (by continent). (C) Bayesian information criteria (BIC) indicating the most probable number of genetic groups. (D) Scatterplot from discriminant analysis of principal components (DAPC).

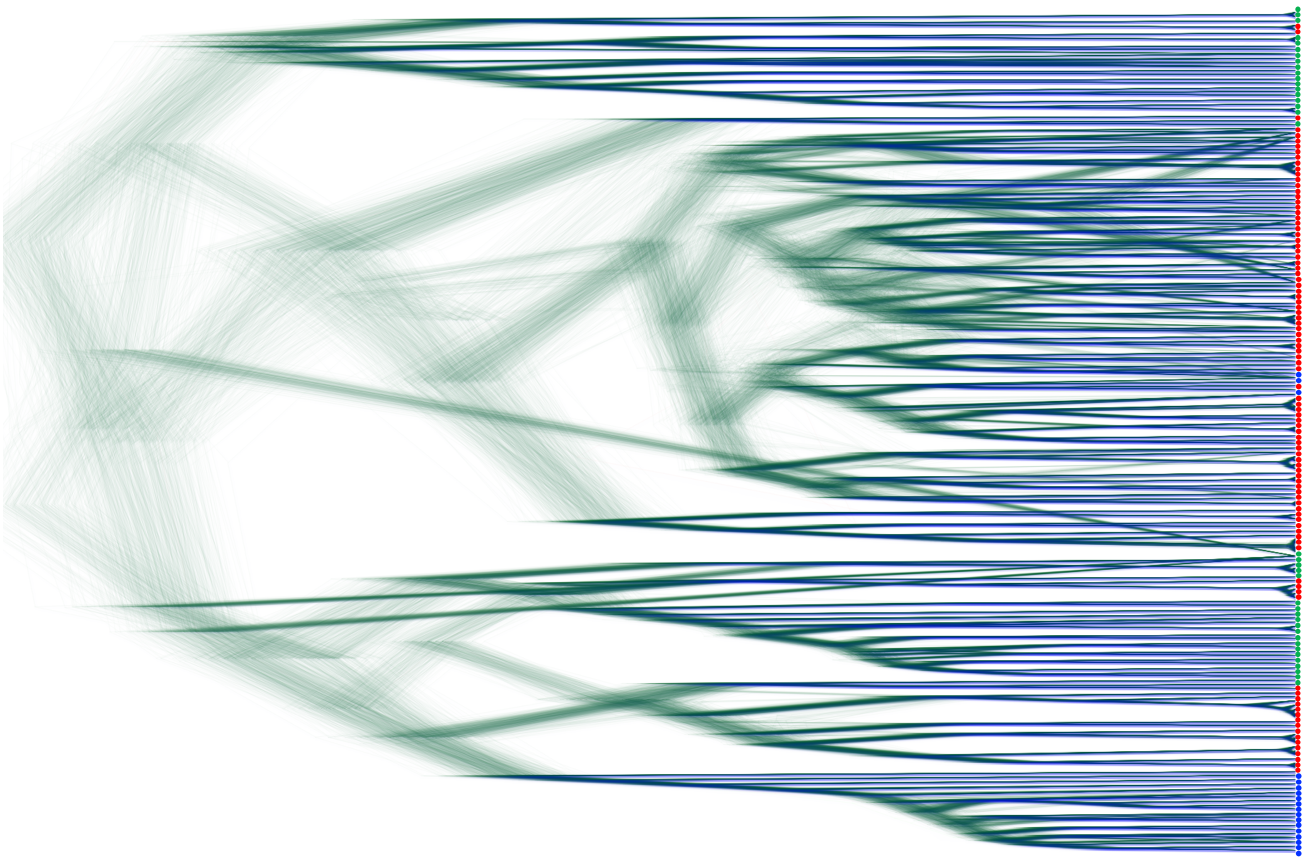

**Figure S5.** Densitree cloudogram based on 203 samples from chromosome 1. Thin lines indicate a possible topology involving one sample. The consensus tree is represented in dark blue. Red dots represent samples from the European lineage (EU), blue dots represent samples from the Brazilian lineage (BR) and green dots represent samples from the North American lineage (NA).

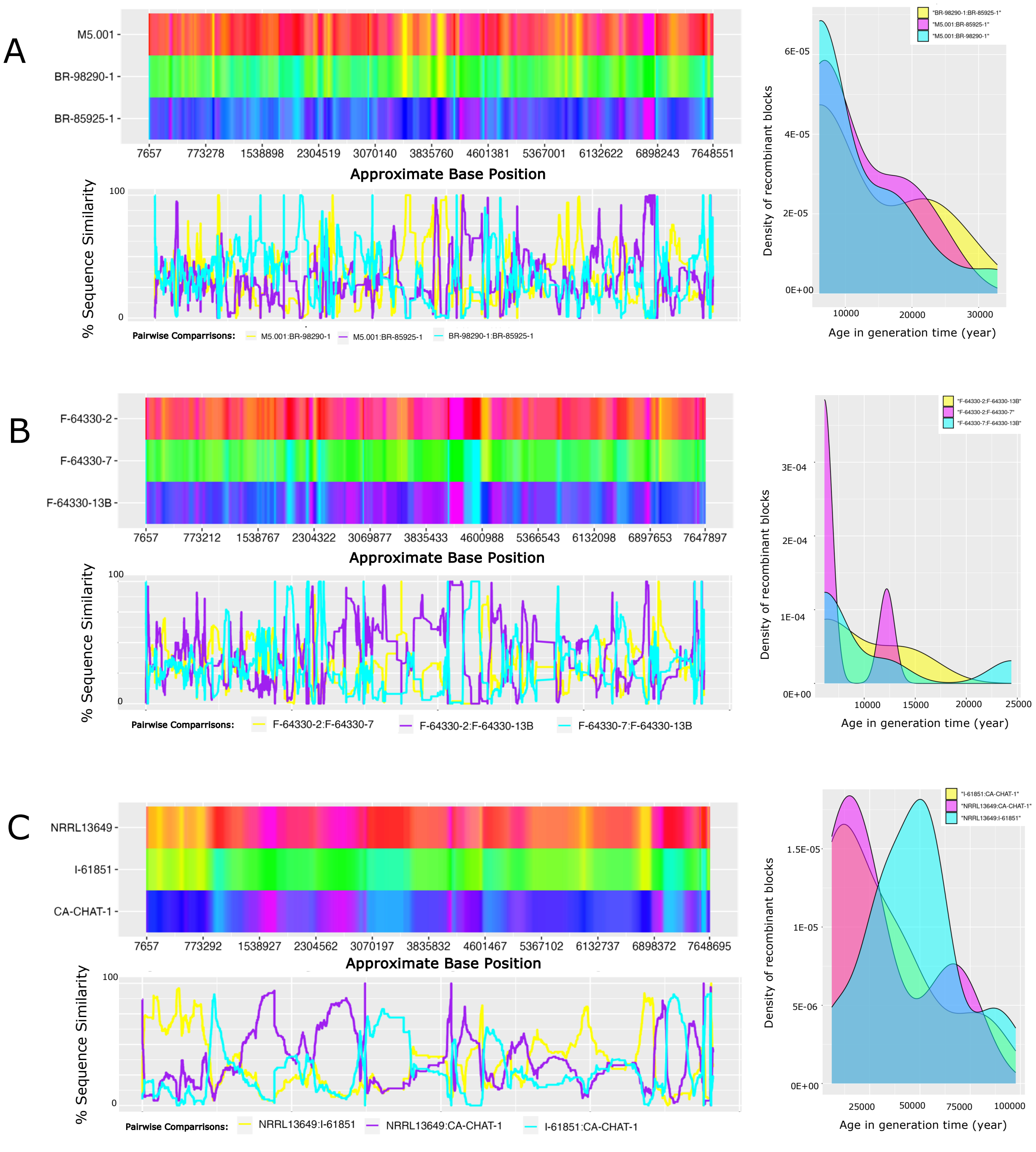


**Figure S6**. Recombination analysis of chromosome 1. (A) Left: Sequence similarity among isolates M5.001:BR-98290-1:BR-85925-1 (triplet 3), visualized through an RBG color triangle by HYBRIDCHECK. Right: Age distribution of recombinant blocks detected by HYBRIDCHECK. (B) Similar analyses using the isolates F-64330-2:F-64330-7:F-64330-13 (triplet 4). (C) Similar analyses using the isolates NRRL13649:I-61851:CA-CHAT-1 (triplet 5).

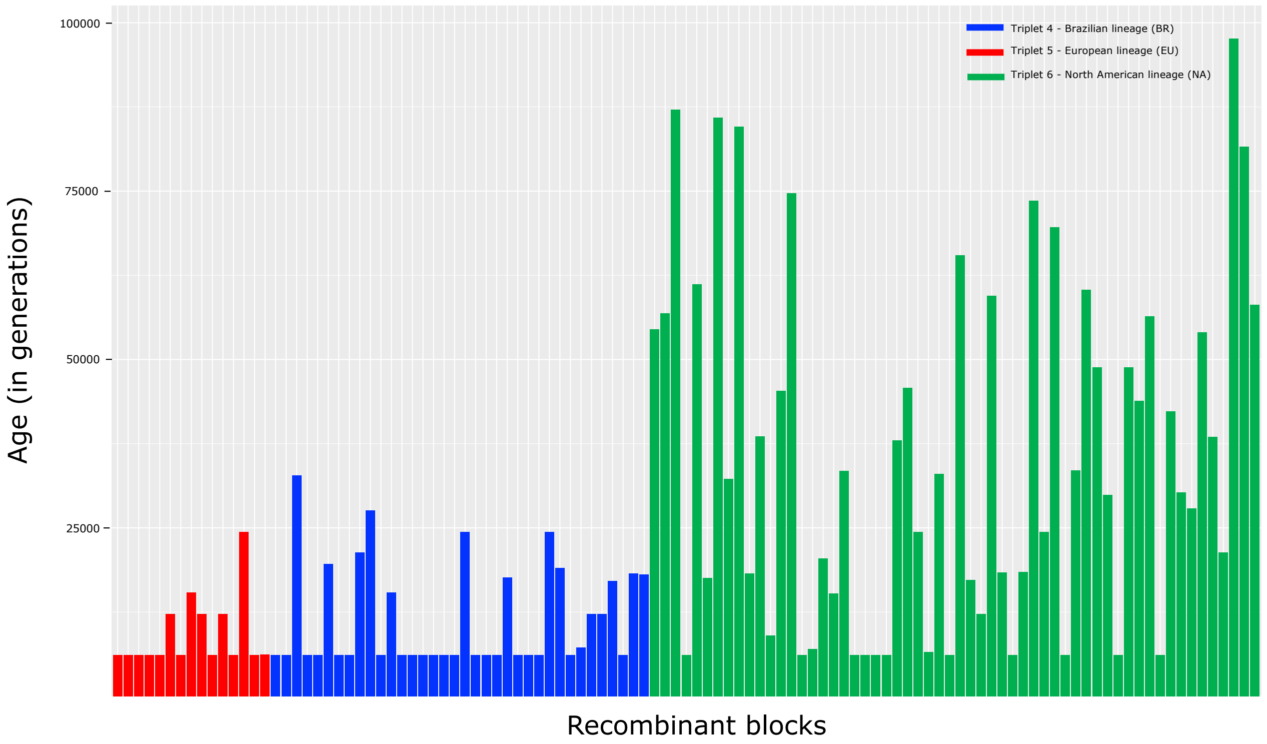


**Figure S7.** Age distribution of recombinant blocks involving Brazilian isolates: M5.001:BR-98290-1:BR-85925-1 (triplet 4), European isolates: F-64330-2:F-64330-7:F-64330-13B (triplet 4), and North American isolates: NRRL13649:I-61851:CA-CHAT-1 (triplet 5). Estimates were generated using HYBRIDCHECK assuming a mutation rate of $10^{-8}$ per generation and a generation time of one year.

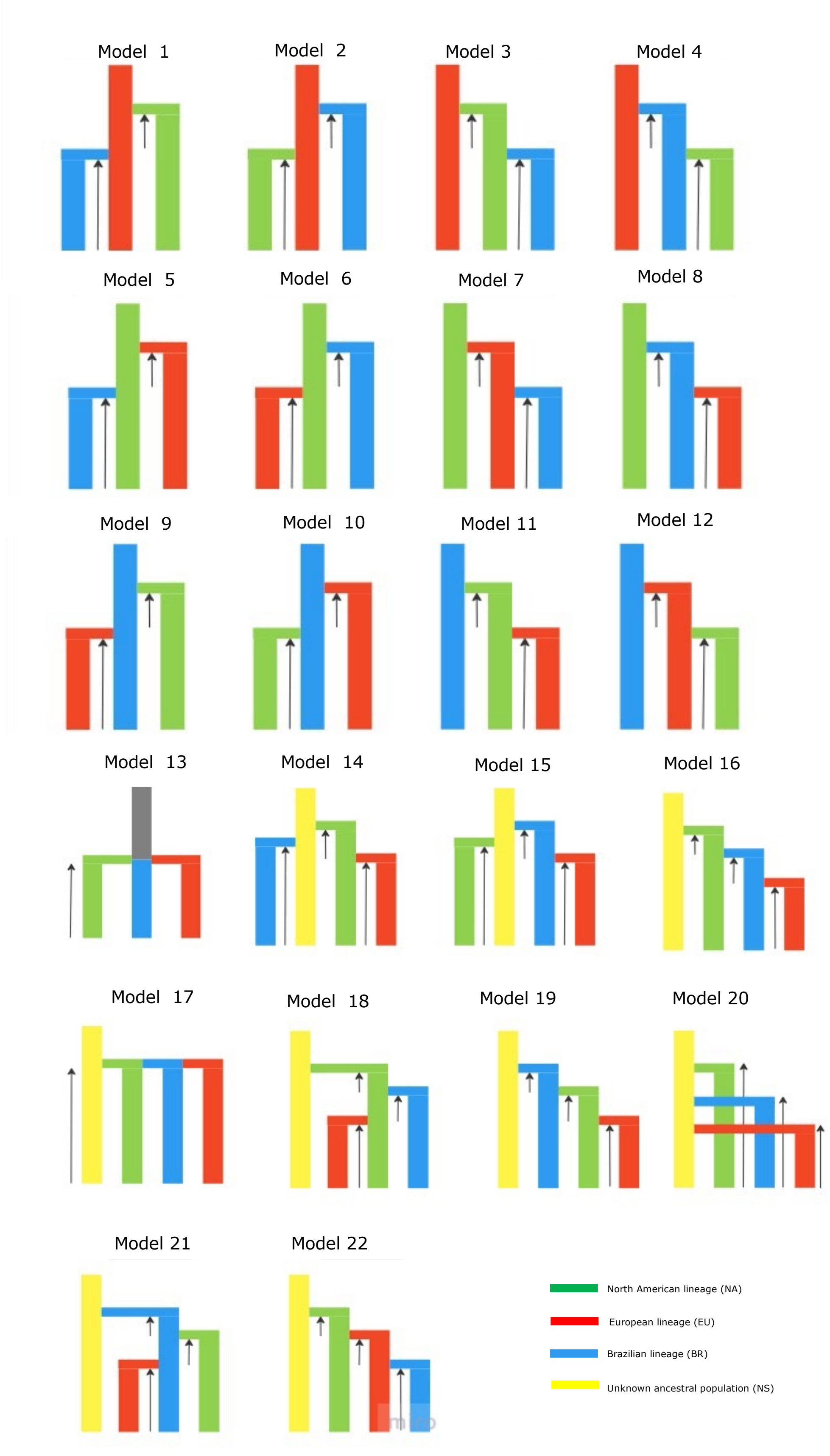


**Figure S8**. Demographic models tested. Colors indicate the genetic lineage of *C. graminicola* and unsampled ancestral lineage. Rows indicate population divergence.

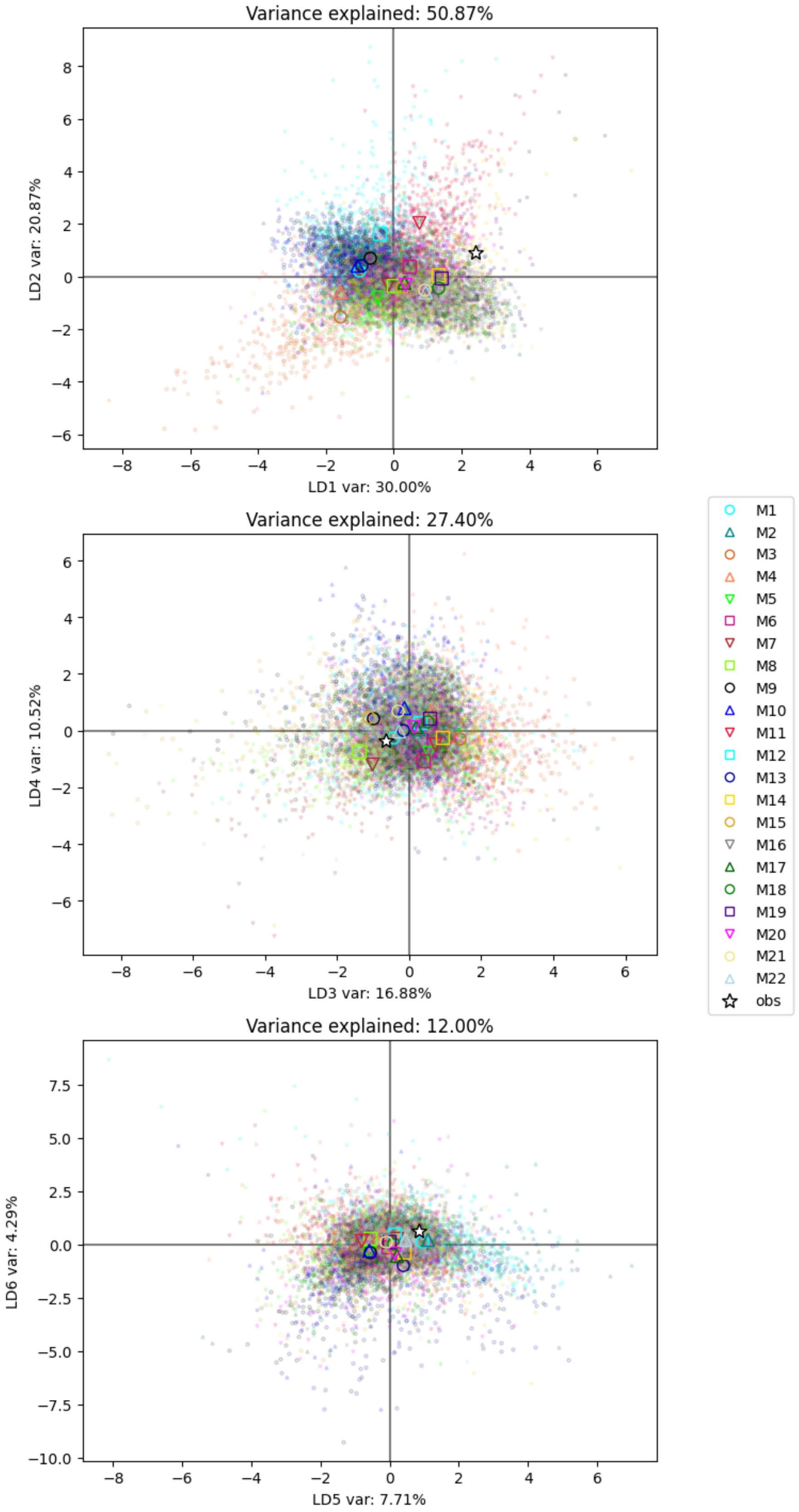


**Figure S9.** The fit of the data used in the ABC analysis of the population history of *Colletotrichum graminicola* using linear discriminant analysis (LDA). Colored dots indicate simulated data from 30 models tested and the star indicates the observed dataset**.**

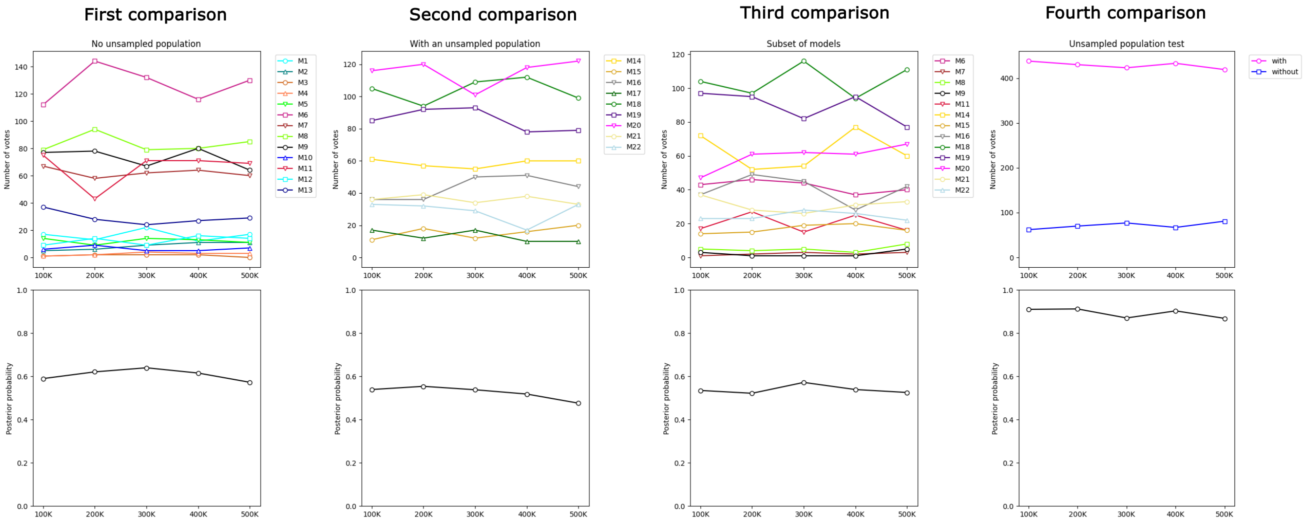


**Figure S10.** Results of model choice analysis. Top: number of votes obtained in the random forest for each model or group of models. Bottom: posterior probability of the best model or group of models. The comparisons are replicated over a range of simulations to assess the consistency of the results.

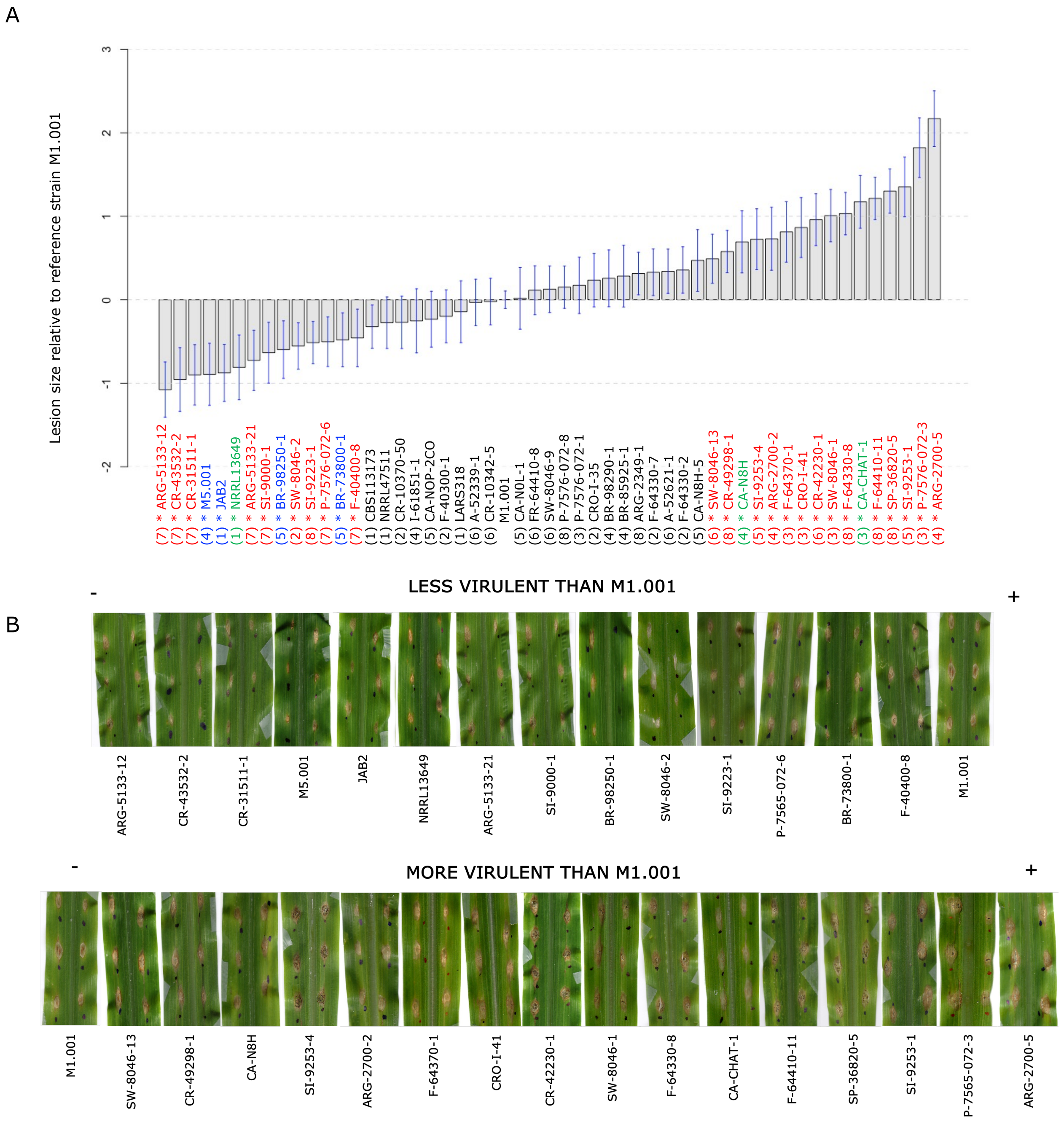


**Figure S11**. Pathogenic characterization of *Colletotrichum graminicola* isolates. (A) Graphic representation of 53 isolates in order of virulence (relative to the reference strain M1.001) in ascending order. Pathogenicity assay batch number is shown in parentheses. A confidence interval using the common estimate of the variability and a Sidäk's correction is shown around the mean virulence of each strain. Two strains are significantly different if their confidence intervals do not have points in common. Isolates marked with an asterisk were significantly different ($P < 0.05$) from strain M1.001 based on a post hoc test using a Sidäk's correction. (B) Necrotic lesions on maize leaves at 4 days after inoculation with spore suspensions of the isolates. Black dots indicate the inoculation points.

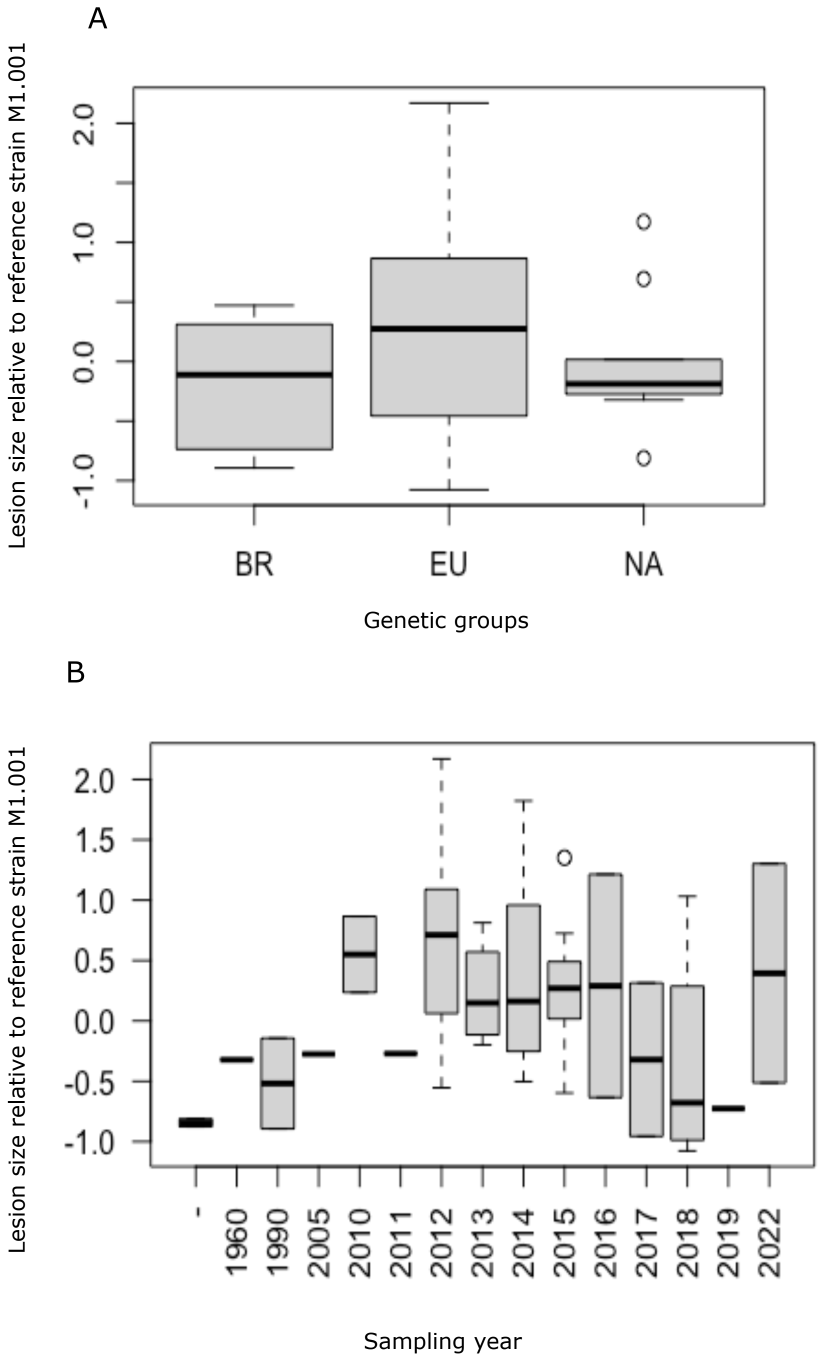


**Figure S12**. (A) Bar plot showing the distribution of the virulence for the three genetic lineages. (B) Bar plot showing the distribution of the virulence when the isolates were grouped by year.
